# Structure-activity relationships of B.1.617 and other SARS-CoV-2 spike variants

**DOI:** 10.1101/2021.09.12.459978

**Authors:** Tzu-Jing Yang, Pei-Yu Yu, Yuan-Chih Chang, Ning-En Chang, Yu-Xi Tsai, Kang-Hao Liang, Piotr Draczkowski, Bertina Lin, Yong-Sheng Wang, Yu-Chun Chien, Kay-Hooi Khoo, Han-Chung Wu, Shang-Te Danny Hsu

**Affiliations:** Institute of Biological Chemistry, Academia Sinica, Taipei, Taiwan; Institute of Biochemical Sciences, National Taiwan University, Taipei, Taiwan; Academia Sinica Cryo-EM Center, Academia Sinica, Taipei, Taiwan; Institute of Cellular and Organismic Biology, Academia Sinica, Taipei, Taiwan; Biomedical Translation Research Center (BioTReC), Academia Sinica, Taipei, Taiwan; Faculty of Pharmacy, Medical University of Lublin, ul. W. Chodzki 4a, 20-093 Lublin, Poland; Johns Hopkins University, Baltimore, MD, U.S.A.

## Abstract

The surge of COVID-19 infection cases is spurred by emerging SARS-CoV-2 variants such as B.1.617. Here we report 38 cryo-EM structures, corresponding to the spike protein of the Beta (B.1.351), Gamma (P.1), Delta (B.1.617.2) and Kappa (B.1.617.1) variants in different functional states with and without its receptor, ACE2. Mutations on the N-terminal domain not only alter the conformation of the highly antigenic supersite of the Delta variant, but also remodel the glycan shield by deleting or adding N-glycans of the Delta and Gamma variants, respectively. Substantially enhanced ACE2 binding was observed for all variants, whose mutations on the receptor binding domain modulate the electrostatics of the binding interfaces. Despite their abilities to escape host immunity, all variants can be potently neutralized by three unique antibodies.

## Introduction

At the end of August 2021, severe acute respiratory syndrome coronavirus 2 (SARS-CoV-2) has infected over 210 million people globally and the resulting coronavirus disease 2019 (COVID-19) has claimed more than 4.4 million lives^1^. Since the identification of the original SARS-CoV-2 strain in Wuhan, China in late 2019, several variants of concern (VOC) with increased transmissibility and reduced sensitivity to antibody neutralization have emerged^2,3^. The Alpha (B.1.1.7) variant was the first VOC, was identified in the United Kingdom in September 2020 and emerged to global dominance that peaked in mid-March 2021. Meanwhile, the Beta (B.1.351) and Gamma (P.1) variants were identified in South Africa (May 2020) and Brazil (November 2020), respectively. Despite their significantly increased transmissibility^4^, resistance to convalescent/vaccination sera and monoclonal neutralizing antibodies (nAbs), and higher likelihood of severe symptoms^5^, their circulation are mostly restricted to their origins and neighboring countries^1^. In early 2021, the B.1.617 lineage was reported in India. It subsequently evolved into the B.1.617.1 and B.1.617.2 sub-lineages that are designated as the Kappa and Delta variants, respectively. Within weeks since its emergence outside India, the Delta variant has become the most dominant VOC, accounting for more than 90% of the reported daily cases to date. In contrast, the Kappa variant does not gain as much prominence and it is currently only designated as a variant of interest (VOI).

The definitions of the Beta, Gamma, Delta and Kappa variants are primarily based on the mutations of the highly glycosylated spike (S) protein that is responsible for host recognition, membrane fusion and viral entry (**Fig. 1A**). The S protein can be divided into the N-terminal S1 subunit and C-terminal S2 subunit, which are responsible for host receptor binding and membrane fusion, respectively^6,7^. The receptor binding domain (RBD) within the S1 subunit binds to the ectodomain of angiotensin converting enzyme 2 (ACE2). The RBD is the most immunogenic domain of the S protein, accounting for up to 90% of the neutralizing antibodies (nAbs) derived from convalescent sera^8-11^. RBD-specific nAbs neutralize SARS-CoV-2 variants by inhibiting ACE2 binding through steric hindrance. A number of nAbs are now in clinical use to treat COVID-19 patients^12^. However, mutations within the RBD can reduce the efficacy of nAb neutralization by disrupting the respective conformational epitopes, such as N501Y in the Alpha variant, E484K in the Beta and Gamma variants, and L452R in the Delta and Kappa variants (**Fig. 1, A** and **B**). The reduced neutralization efficacy associated with these mutations indeed lead to the Food and Drug Administration of the United States to revoke the emergency use authorization of bamlanivimab (LY-CoV555)^13^. The S1 subunit, also referred to as the N-terminal domain (NTD), harbors eight N-glycosylation sites that form a glycan shield over the protein surface rendering it less immunogenic^14-16^. Nevertheless, a supersite within the glycan shield has been identified to be targeted by a panel of nAbs^17^. The mechanism by which NTD-specific nAbs neutralize SARS-CoV-2 infection is not well understood; however, it is well established that several VOC- and VOI-associated NTD mutations significantly reduce or completely abolish nAb neutralization thus leading to immunity escape^16,18-20^. Specifically, a recent study reports complete escape of the Delta variant from a panel of 16 NTD-specific nAbs^21^.

**Fig 1.**
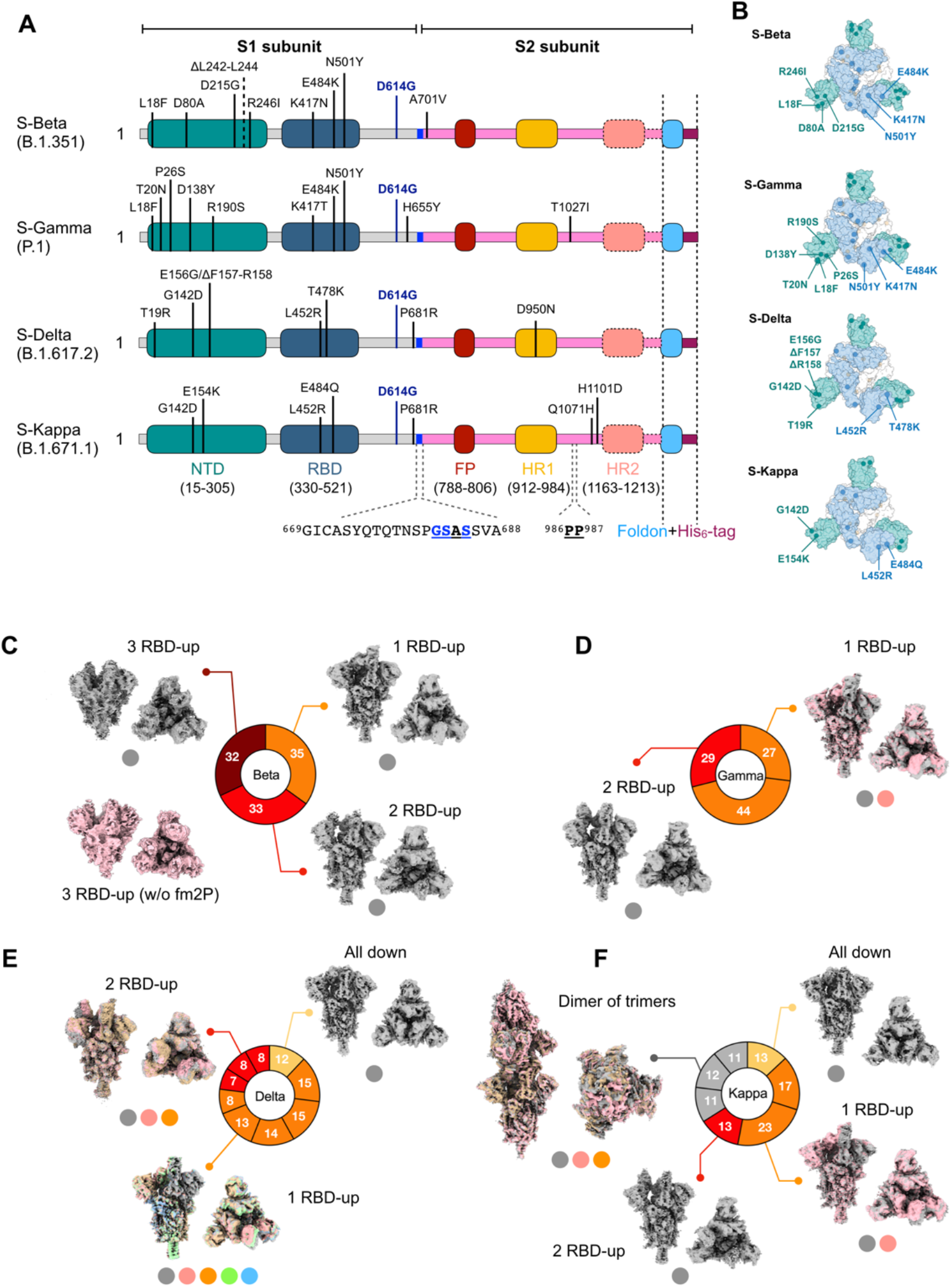
Conformational landscape of the spikes of the SARS-CoV-2 Beta, Gamma, Delta and Kappa variants. (A) Domain organization and mutations of SARS-CoV-2 variants. (B) Spatial distribution of the mutations on the S variants. The representative cryo-EM structures are shown in semitransparent surface with the mutation sites indicated in spheres with their identities labeled. (C-F) S protein conformational distributions categorized according to the RBD-up state as indicated in the pie charts. Zero, one, two and three RBD-up states are colored in yellow, orange, red and brick, respectively. Within each basin of the RBD-up state, contributions of individual sub-states are indicated by the solid lines. The unique dimer of S-Kappa trimer is colored in grey in the pie chart. Orthogonal views the cryo-EM maps of individual RBD-up states are shown around the pie charts. In cases where multiple substates were observed, they are superimposed and colored as indicated by the filled circles below. The default color is grey, followed by pink, lime, pale blue and orange. S-Delta exhibited the most conformational diversity within the 1 RBD-up and 2 RBD-up states. In particular, the 1 RBD-up state of S-Delta can be classified into five distinct sub-states with clear structural difference not only within the RBD but also in the NTD and the HR1.

As global vaccination programs have expanded at an unprecedented pace, reductions in the effectiveness of vaccines against different VOCs are emerging. For instance, the Pfizer/BioNTech (BNT162b2) and Moderna (mRNA-1273) vaccines experience an astounding drop in protection (up to 12-fold) against the Beta variant while their protection against the Gamma variant is reduced by 2-to 3-fold^15,20,22^. While mRNA-1273 affords comparable protection against the Alpha variant compared to the wild-type (WT) Wuhan strain^20,22-24^, a 2-to 3-fold reduction in protection against the Delta variant is reported^25,26^. In contrast, BNT162b2 generates substantially less neutralizing titers against the Delta variant compared to the Alpha variant^21,27^. Another study shows that the Kappa variant is even more resistant to BNT162b2-induced sera compared to the Delta variant and other emerging variants^28^. Notably, one study reports that the Delta and Beta variants essentially abolish the neutralizing activity of the AstraZeneca (ChAdOx1 nCoV-19) vaccine^29^. Notably, convalescent sera from the Beta or Gamma variant-infected individuals and the vaccine-elicited antibodies show substantially reduced neutralization against the Delta and Kappa variants^11,30^. These findings raise the questions about of the current vaccines against emerging breakthrough SARS-CoV-2 variants^31^. As current developments of COVID-19 vaccines are chiefly focused on the S protein, understanding how the mutations associated with VOCs and VOIs on the structure and function of the S protein, and their interactions with ACE2 and nAbs, is therefore of paramount importance.

## Results

### Conformational landscape of SARS-CoV-2 spike variants

To examine the effect of mutations on the structures of the S protein of the Beta (S-Beta), Gamma (S-Gamma), Delta (S-Delta) and Kappa (S-Kappa) variants, we solved in total 25 cryo-EM structures of the apo S protein variants, revealing a vast conformational landscape for each of the variants (**Fig. 1C-F** and **Extended Data Figs. 1-5**). S-Beta exhibited a highly populated fully open state, i.e., all three RBDs are in the up conformation (3 RBD-up) in addition to the partially open states in which one or two RBDs were in the up conformation (1 RBD-up or 2 RBD-up; **Fig. 1C** and **Extended Data Fig. 2, Extended Data Table 1** and **Extended Data Movie 1**). To verify the authenticity of the unique 3 RBD-up conformation, we solved the cryo-EM structure of S-Beta without the stabilizing tandem proline and furin cleavage site substitutions (fm2P; see Methods). A well-defined 3 RBD-up state could also be identified for the cleavable form of S-Beta (**Extended Data Fig. 2** and **Extended Data Table 2**). While S-Gamma only populated the 1 RBD-up and 2 RBD-up states, the 1 RBD-up state of S-Gamma could be divided into two distinct substates through a robust three-dimensional variability analysis (3DVA; **Fig. 1D, Extended Data Fig. 3, Extended Data Table 3** and **Extended Data Movie 2**). Of the four S variants under investigation, S-Delta exhibited the largest conformational diversity, encompassing five substates within the 1 RBD-up conformational basin and three substates within the 2 RBD-up conformational basin, in addition to the fully closed conformation of which all three RBDs are in the down conformation (all RBD down) that was hitherto not observed in the D614G and Alpha (B.1.1.7) S variants (**Fig. 1E, Extended Data Fig. 4, Extended Data Table 4** and **Extended Data Movie 3**)^32,33^. The separation of multiple substates within the same class (conformational basin) of the RBD-up conformation was made possible through two rounds of 3DVA (**Extended Data Fig. 4** and Methods). Unexpectedly, we identified an unprecedented head-to-head dimer of S-Kappa trimers, involving all three RBDs in each trimeric S protein to bind to another RBDs from the other trimeric S protein. Furthermore, S-Kappa exhibited an all RBD down conformation, a 1 RBD-up conformation with two substates, and a single 2 RBD-up conformation (**Fig. 1F, Extended Data Fig. 5, Extended Data Tables 5-6**, and **Extended Data Movie 4**). The head-to-head dimer of S-Kappa trimers displayed rigid body motions between the two trimers that could be resolved into three substates. Collectively, our comprehensive structural analyses of the SARS-CoV-2 S variants illustrates a broad conformational space explored by each of the variants with S-Delta and S-Kappa being the most intriguing in terms of the large number of substates within each basin of the RBD conformational grouping, and the unique higher-order assembly, respectively.

### Structural basis of immunity escape of S-Delta and S-Gamma

Being the most prevalent COVID-19 VOC to date with increased transmissibility and immunity escape, the Delta variant harbors ten mutations in the S protein compared to the original Wuhan strain (S-WT), namely T19R, G142D, E156G, Δ157, Δ158, L452R, T478K, D614G, P681R and D950N^27^ (**Fig. 1, A** and **B**). Recent studies show that VOCs are refractory to vaccine and convalescent sera as well as nAbs that target the RBD and NTD to different degrees^21,27,30,34^. The Delta variant, in particular, shows reduced sensitivity to convalescent sera from the Beta and Gamma variants, indicating that the Beta, Gamma and Delta variants are antigenically divergent, and that there is a risk of reinfection by the Delta variant for individuals who have been infected by the Beta or Gamma variants^30^. The Delta variant is also completely refractory to NTD-specific nAbs isolated from convalescent sera with prior infection of the WT strain^21^. As the mutations within the NTD are the most divergent among the VOCs and VOIs, we compared the structure of the NTD of S-Delta with respect to that of S-WT. Structural mapping of the epitope count based on the reported structures of NTD in complex with antibodies revealed a supersite encompassing a cluster of residues that are most antigenic, including Y145, H146, L249 and P251 (**Fig. 2A** and **Extended Data Table 7**)^35^. The double deletions (Δ157 and Δ158) and the substitutions, G142D and E156G, resulted in major conformational rearrangements of the supersite, including the sequestering of Y145 (**Fig. 2B**). The loop on which the N149 glycan is located underwent significant conformational rearrangements that greatly alters the glycan shield around the supersite (**Fig. 2B** and **Extended Data Fig. 6A**). Furthermore, the T19R mutation abrogates the ^17^NLT^19^ sequon for N-glycosylation. Indeed, we did not observe any N-glycan density in the cryo-EM maps of S-Delta (**Extended Data Fig. 6A**) nor did we find evidence of glycosylation in mass spectrometry (MS) analysis (**Extended Data Fig. 7**).

**Fig 2.**
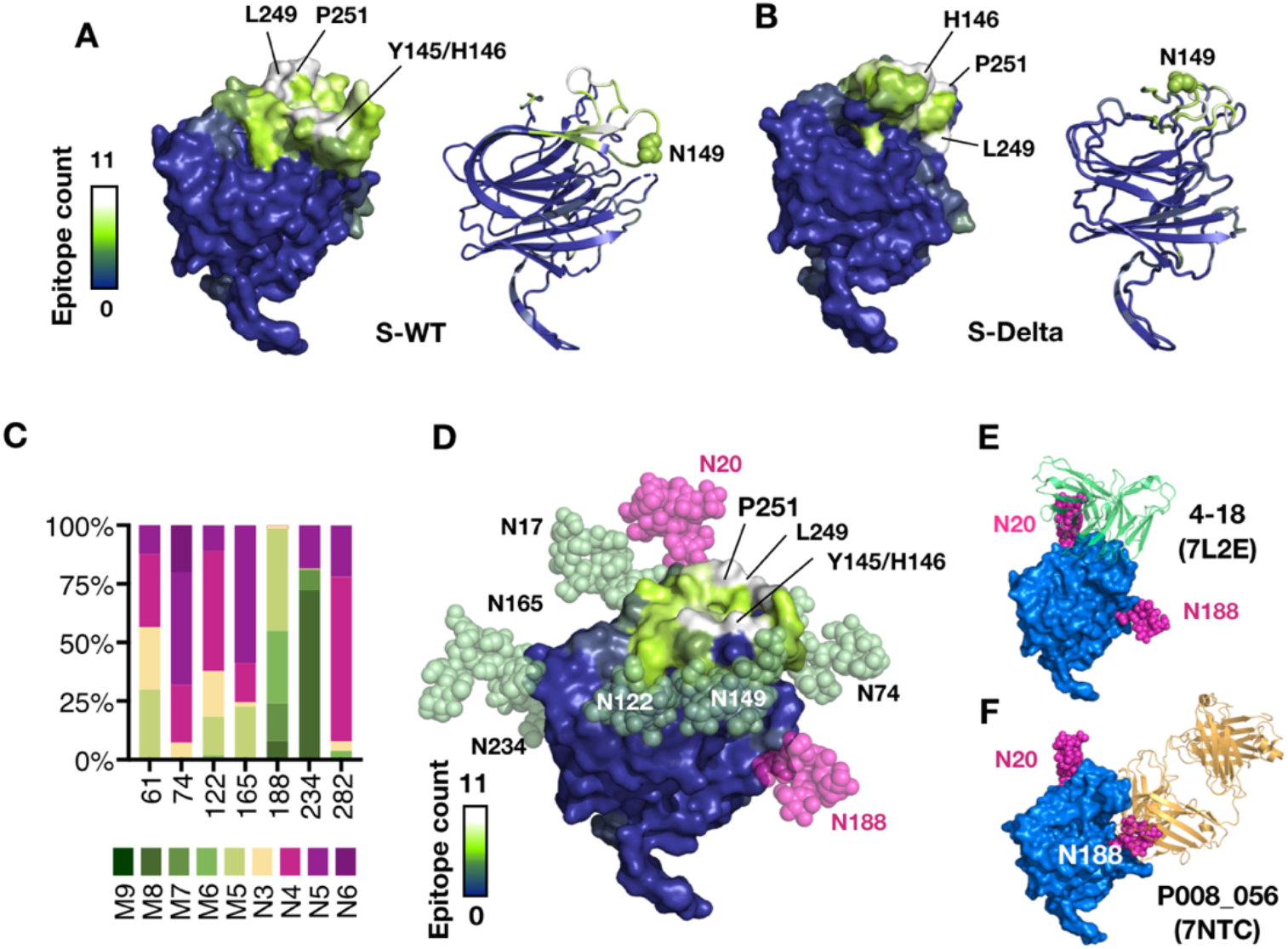
Impacts of the mutations on the glycosylation around the NTD supersite of S-Gamma. Surface and cartoon representations of the NTD of (A) S-WT and (B) S-Delta color-coded by the epitope count as indicated on the left. The most antigentic residues are indicated in the surface representation. N149 that is N-glycosylated is shown in green spheres. (C) MS-based quantitation of the glycoform distributions of individual N-glycosylation sites within the NTD of S-Gamma. The contributions of different glycoforms are color-coded according to the labels below, which are sorted from left to right, corresponding to degree of processing during the post-translational modification of which Man9 is the most under-processed glycoform. (D) Atomic model of the fully glycosylated NTD of S-Gamma based on the cryo-EM and MS analyses. The protein part is presented in the same scheme as in (A). The N-glycans are shown in semitransparent green spheres except for N20 and N188, shown in magenta to indicate their unique steric contributions occlude nAb binding as shown in (E) and (F) for nAb 4-18 and P008_056, respectively. The nAb-bound NTD is shown in blue surface with the N20 and N188 glycans manually docked onto the respective side chains. Fab of 4-18 and P008_056 are shown in semitransparent lime ad gold cartoon representation, respectively. The PDB codes are shown in parentheses.

In the context of immunity evasion through alteration of the glycan shield, S-Gamma harbors two NTD mutations – T20N and R190S – that introduce two additional N-glycans on N20 and N188. MS analysis unambiguously established the presence of a high-mannose type N-glycan at N188 with five mannose residues (M5) being the predominant glycoform (**Fig. 2C**). The remaining N-glycans on the NTD are primarily complex types, except for the N234 glycan, which is mostly high-mannose type, in line with previous reports^14^. We note that a shift from mannose-9 (M9) to mannose-8 (M8) together with a small proportion of highly processed complex type glycan (N5) were observed for N234, suggesting that the N234 glycan is less occluded during Golgi glycosylation compared to other spike variants that show exclusively a high-mannose type glycan at N234 with the major glycoform being M8^14^ (**Extended Data Fig. 7**). The cryo-EM map of S-Gamma showed additional protrusion from the side-chain of N20 indicative of the presence of N-glycan (**Extended Data Fig. 6B**). Molecular modeling of a fully glycosylated NTD of S-Gamma showed that the supersite is surrounded by N-glycans (**Fig. 2D**), and that the additional N20 and N188 glycans can sterically hinder NTD-specific antibody binding such as nAb 4-18, which targets the supersite^19^ (**Fig. 2E**), and nAb P008_056, which targets an alternative epitope, requiring major conformational rearrangements in the loops on which N188 resides^18^ (**Fig. 2F**). These findings provided a clear structural basis underlying the immunity escape of the Gamma variant through the NTD mutations.

### Effect of mutation on the higher order assembly of S-Kappa

Our cryo-EM analysis showed that S-Kappa exhibited an ensemble of conformations with one third adopting an unprecedented head-to-head dimer of trimers (**Fig. 1F, Fig. 3A** and **Extended Data Movie 4**). The dimeric interface involves (i) inter-RBD van der Waals contacts between F490 and the S-Kappa-specific substitution E484Q, (ii) a bipartite hydrogen bonding network formed between the side chains of Q493 from the dimerizing RBDs, (iii) a bipartite hydrogen bonding network between the side-chain of N450 from one RBD to the backbone amide and side chain carboxyl group of N487 from the other RBD, and (iv) a hydrogen bond between the side chain hydroxyl group of Y489 of one RBD and side chain carboxyl of N450 from the other RBD (**Fig. 3B**). While the N450-mediated hydrogen bonding is asymmetrically formed between N487 or Y489 from the other RBD, the van der Waals contacts and the bipartite hydrogen bonding associated with Q493 are stably formed in all three pairs of RBDs. The higher order assembly of S-Kappa can be attributed to the unique E484Q substitution that is primarily found in the Kappa (B.1.617.1), B.1.617.3, B.1.630 and P5 variants per PANGO lineage but not in other VOCs and VOIs^1^. The negatively charged E484 in S-WT and S-Delta will create unfavorable electrostatic repulsion when positioned in the same higher-order arrangement. Likewise, the positively charged E484K substitution in S-Beta and S-Gamma will generate the same electrostatic repulsion to prevent the dimer formation.

**Fig 3.**
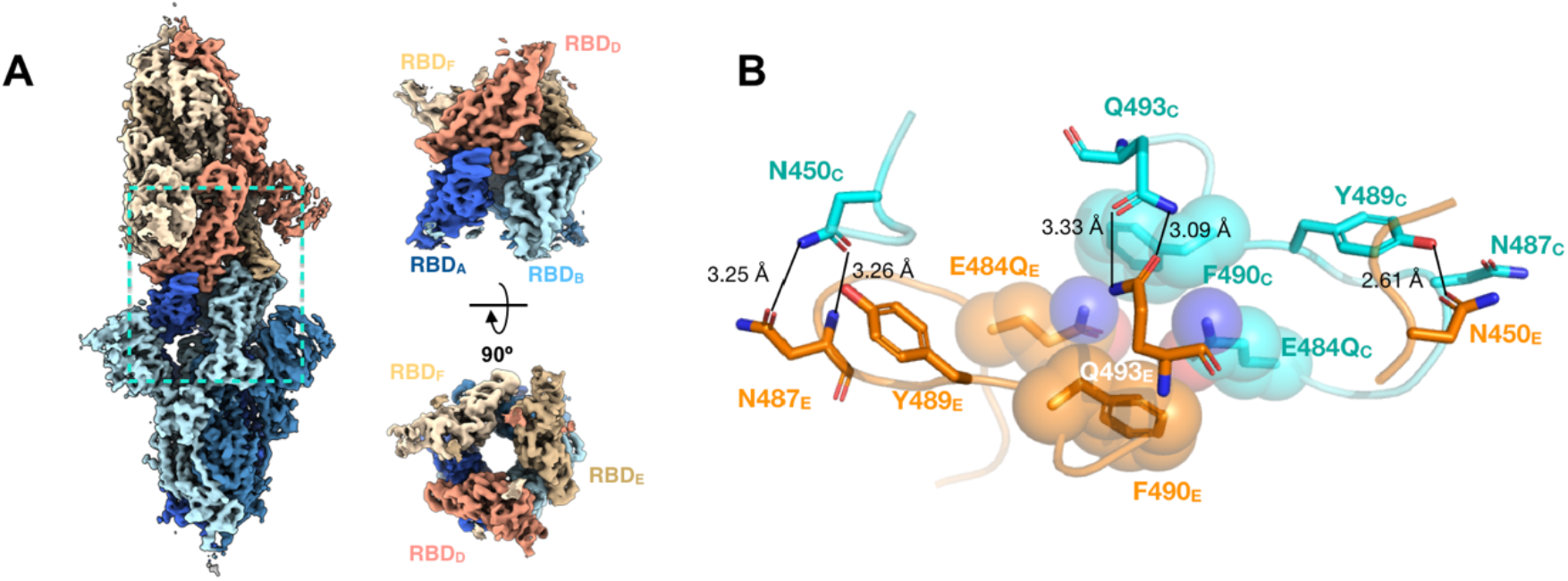
S-Kappa forms a unique head-to-head dimer of trimers. (A) A representative Cryo-EM map of the dimer of S trimers with one trimeric S-Kappa shown in differ shades of blue and the other shown in different shades of orange. The dimer interface involving the RBDs is boxed in a dashed blue rectangle, which is expanded and shown in orthogonal views on the right. (B) Detailed dimer interface interactions of a pair of RBD. The interacting residues are shown stick representation with the nitrogen and oxygen atoms shown in blue and red, respectively. The carbon atoms are colored in light blue and orange for chain C and E, respectively. The residues involved in van der Walls contact are shown in semitransparent spheres. Inter-molecular hydrogen bonds are shown in black line with the bond distance indicated.

### Structural basis of enhanced receptor ACE2 binding of spike variants

We next investigated the effects of mutations on the receptor ACE2 binding by biolayer interferometry (BLI). Superfold GFP-fused ACE2 ectodomain (hereafter ACE2) was immobilized on the BLI sensor to bind to different concentrations of S-Beta, S-Gamma, S-Delta and S-Kappa, all of which showed significantly enhanced ACE2 binding with respect to that of S-D614G with an increase in the dissociation constant (K_d_) by 12.4, 10.3, 7.9 and 34.4-fold, respectively (**Fig. 4A** and **Extended Data Table 8**). Among the four S variants, S-Kappa bound to ACE2 the strongest, with a K_d_ of 0.09 nM. To glean structural insights into the effect of mutations on the enhanced receptor binding, we determined 13 cryo-EM structures of the S variants in complex with ACE2, aided by local refinements masked around RBD and ACE2 to improve the resolution of the ACE2-binding interface (**Fig. 4B, Extended Data Figs. 8-11** and **Extended Data Tables 9-12**). Using the aforementioned 3DVA procedure, we determined a cryo-EM structure of S-Beta in complex with ACE2 in a 3:3 binding stoichiometry (3 ACE2-bound; **Fig. 4C**). On the one hand, a minor fraction of S-Gamma, S-Delta, and S-Kappa was found to bind to ACE2 in a 3:2 stoichiometry (2 ACE2-bound). On the other hand, two distinct 3 ACE2-bound substates were observed for S-Gamma and S-Delta. By using the most representative ACE2-RBD structure of each S variant to determine the changes of solvent accessible surface area (ΔSASA), we found a strong correlation between the free energy derived from the K_d_ values (ΔG = –RT ln K_d_) and ΔSASA (**Fig. 4D**). Close examination of the binding interface between RBD and ACE2 showed highly complementary surface electrostatic potentials with RBD being positively charged and ACE2 being negatively charged (**Fig. 4E** and **Extended Data Fig. 12**). The surface electrostatic distribution on the RBD was clearly modulated by the mutations in different variants. K417N in S-Beta and K417T in S-Gamma reduce the positive charge; E484K in S-Beta and S-Gamma increase the positive charge. Similarly, T478K in S-Delta, and L452R in S-Delta and S-Kappa increase the positive charge. E484Q, which is responsible for the dimerization of S-Kappa trimer, reduces the negative charge to favor the electrostatics-driven ACE2 binding (**Fig. 4E** and **Extended Data Fig. 12**). Collectively, the mutations in S-Kappa considerably reshape the positive isoelectrostatic potential around the ACE2 binding residues centered around N501. Although the N501Y substitution makes a favorable π-π stacking to enhance ACE2 binding in S-Alpha, the enhancement in K_d_ was only 2.3-fold according to our previous study under the same condition as opposed to the remarkable 34.4-fold increase for S-Kappa.

**Fig 4.**
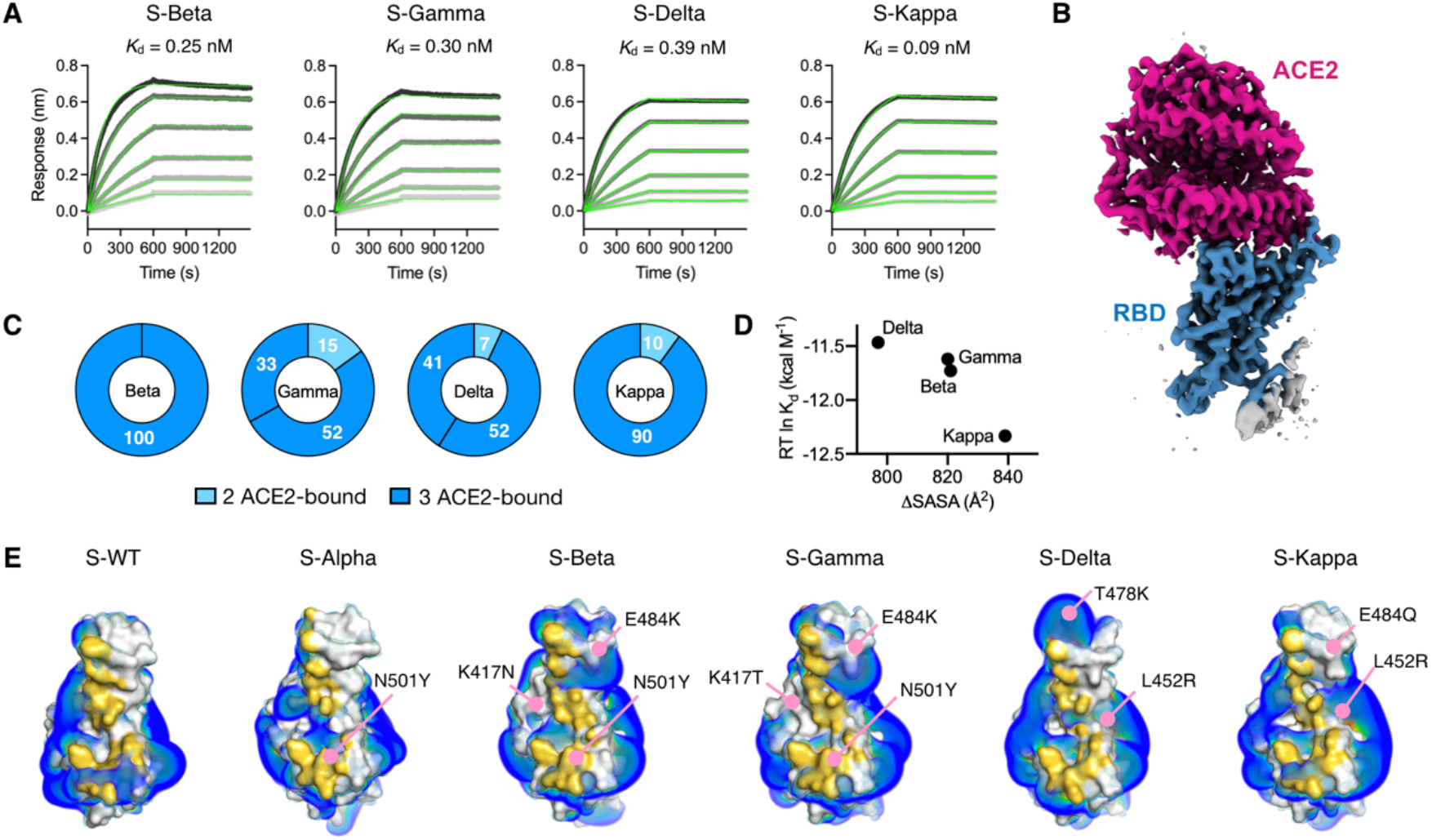
Kinetics and structural basis of receptor ACE2 binding by the SARS-CoV-2 variants. (A) BLI sensorgrams of immobilized ACE2 binding to the S variants. The sensorgrams of the 100, 50, 25, 12.5, 6.25 and 3.125 nM S variants are colored from black to light grey in descending orders superimposed with the fitting results in green. The corresponding Kd values are shown above. (B) Representative cryo-EM map of the RBD S-Delta in complex with ACE2 is colored in blue and magenta for RBD and ACE2, respectively. (C) Distributions of distinct receptor ACE2 binding stoichiometry and quaternary conformations classified according to the number of RBD-bound ACE2. The coloring scheme is shown below. (D) Correlation between the free energy of ACE2 binding and ΔSASA. (E) Surface representations of the RBD of the individual variants with the binding interfaces of the S variants highlighted in gold. Positive isosurfaces of the RBD of the S variants were generated by the APBS module within PyMol. The positions of individual mutations are indicated by pink lines with their changes indicated. The structure of S-WT RBD in complex with ACE2 is taken from the PDB ID 6M0J. The structure of S-Alpha RBD in complex with ACE2 is taken from the PDB ID 7EDJ.

### Broad-spectrum neutralization of antibodies against spike variants

In light of the increased risk of immunity escape by the SARS-CoV-2 variants, antibodies that exhibit broad-spectrum neutralization activities are imperative^34,35^. We previously reported three potent RBD-specific monoclonal nAbs against WT, D614G and Alpha variants, namely RBD-chAb-15, −25 and −45, which have non-overlapping structural epitopes^32^. The neutralizing activity of RBD-chAb-25 against the Alpha variant is lost due to the N501Y mutation while a cocktail of RBD-chAB-15 and −45 can effectively neutralize the D614G and Alpha variants with a half-maximum inhibitory concentration (IC_50_) value of 0.11 and 1.18 ng ml^-1^, respectively. As the Beta and Gamma variants also harbor the N501Y mutation (**Fig. 1, A and B**), RBD-chAb-25 could not effectively compete with ACE2 binding (**Fig. 5A**). Nevertheless, RBD-chAb-25 remained highly effective in competing ACE2 binding against S-Delta and S-Kappa. Importantly, RBD-chAb-15 and −45 robustly inhibited ACE2 binding for all four S variants. Furthermore, negative staining electron microscopy (NSEM)-based structural analyses revealed that the cocktail of RBD-chAb-15 and −45 bound to all four S variants with a 3:3:3 stoichiometry in the same poses as previously reported for S-D614G (**Fig. 5B, Extended Data Fig. 13** and **Extended Data Table 13**). The atomic structure of RBD in complex with BD-chAb-15 and −45 fit well with the EM maps of all four S variants bound to the antibody cocktail (**Fig. 5B** and **Extended Data Fig. 13**). The same analysis showed that RBD-chAb-25 bound to S-Delta in a 3:3 binding stoichiometry in the same pose as previously reported for S-WT; in contrast, the NSEM-derived EM maps showed only one well-defined nAb on the S-Kappa trimer while a significant fraction of S-Kappa was free, and in an all RBD-down form (**Fig. 5B** and **Extended Data Fig. 13**). Of the three RBD-specific nAbs, RBD-chAb-45 was the most potent in neutralizing the all four variants of the SARS-CoV-2 pseudovirus with an IC_50_ value of 3.46, 0.47, 20.1, and 6.68 ng ml^-1^ for the Beta, Gamma, Delta and Kappa variants, respectively (**Fig. 5C** and **Extended Data Table 14**). While RBD-chAb-15 could also neutralize all four pseudovirus variants, the corresponding IC50 values increased significantly to 51.6, 6.03, 125 and 123 ng ml^-1^, respectively. RBD-chAb-25 neutralized the Delta and Kappa variants with an IC50 value of 47.7 and 57.5 ng ml^-1^, respectively. In summary, regardless of the mutations on the RBD that render the individual SARS-CoV-2 variants refractory to many nAbs, including some that are in clinical use, our RBD-specific nAbs displayed broad neutralizing activities against all the emerging SARS-CoV-2 variants with the exception of RBD-chAb-25, which is sensitive to the N501Y mutation present in the Alpha, Beta and Gamma variants.

**Fig 5.**
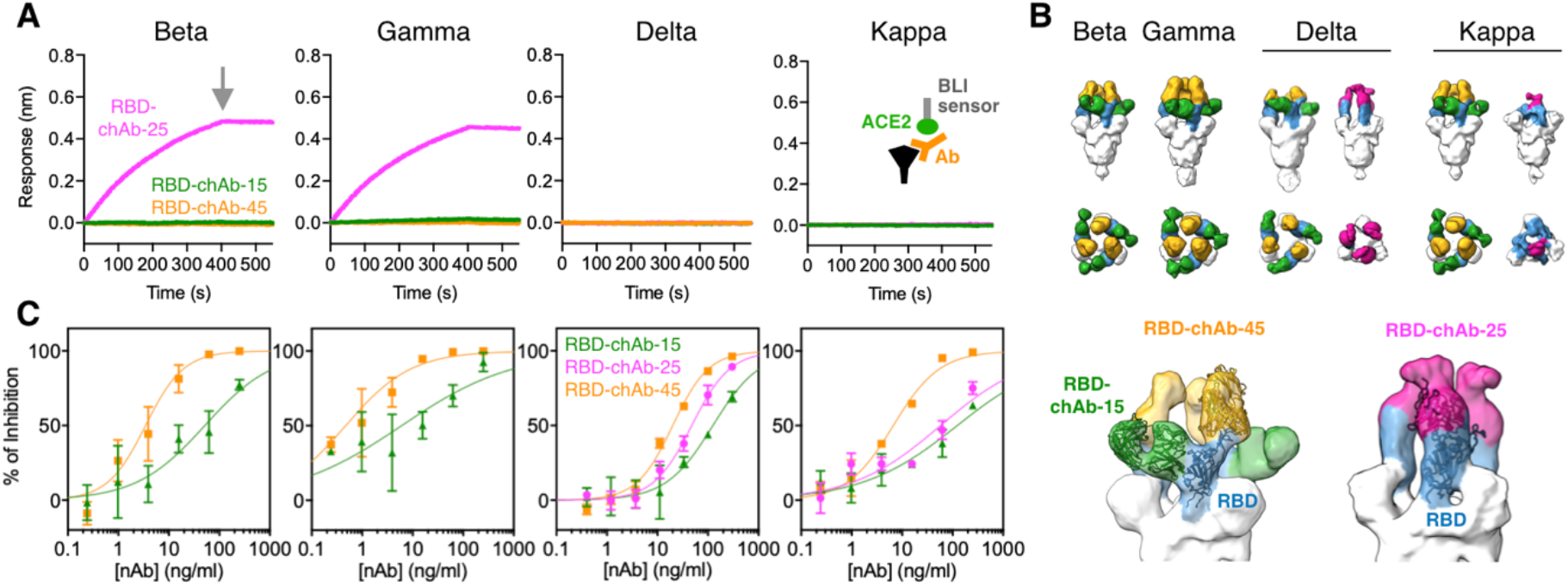
Neutralization of receptor ACE2 binding and pseudovirus infection by three potent nAbs. (A) Competitive ACE2 binding analysis by BLI. RBD-chAb-15, −25 and −45. The invariability of the BLI sensorgrams indicates the effective inhibitation of ACE2 binding by the nAbs as illustrated schematically in the inset of S-Kappa. All nAbs effectively neutralize the ACE2 binding, except for RBD-chAb-25 (magenta), which failed to neutralize the ACE2 binding to S-Beta and S-Gamma. The grey arrow indicates the time point at which dissociation was triggered. (B) NSEM structures of the S variants in complex with the cocktail of RBD-chAb-15/45 or RBD-chAb-25 alone. Top panel. Orthogonal views of the NSEM maps in which the S variants are colored in white, the RBD in blue, and RBD-chAb-15, −25 and −45 in green, magenta and gold, respectively. Bottom panel. Expanded views of RBD-chAb-15/45-bound S-Beta (left) and RBD-chAb-25-bound S-Delta (right). The NSEM maps are shown in semitransparent surface superimposed with the previously reported atomic models: PDB ID 7EH5 for the nAb cocktail and PDB ID 7EJ4 for the RBD-chAb-25. (C) nAb neutralization against different SARS-CoV-2 pseudovirus variants. The data points correspond the mean values of three technical replicates and the error bars correspond to the standard deviation. The solid lines correspond to the fitting result to derive the half-maximum inhibitory concentration (IC_50_) values of individual nAbs (Methods).

## Discussion

In light of the SARS-CoV-2 Delta variant sweeping the globe, and the risk of breakthrough infections, even after vaccination or prior infection by the other SARS-CoV-2 variants^15,20,21,27,30,31,34,35^, our findings provide critical molecular insights into the basis of enhanced receptor binding, host immunity escape and potential treatments based on our potent nAbs not only for the Delta variant but also to the Beta, Gamma and Kappa variants. Efficient host receptor ACE2 binding requires the RBD in an open, upward conformation. We demonstrated that the four S variants displayed a broad range of intrinsic RBD-up propensities with different numbers of RBD-up and multiple substates within the basin of a given RBD-up conformation, exemplified by the five substates in the 1 RBD-up and three substates in the 2 RBD-up conformations for S-Delta (**Fig. 1E** and **Extended Data Movie 3**). The conformational diversity of the RBD of S-Delta is the largest that has been documented to date. A recent study on the full-length S-Delta reports a single all RBD-down and two 1 RBD-up conformations without observing the 2 RBD-up conformations.^36^ The other study on S-Delta focuses on the binding by the NTD- and RBD-specific nAbs.^37^ In this case, a highly engineered HexaPro variant is used, which may alter the propensity of RBD up/down conformations. The common finding from these studies, including ours, is the reorganization of the NTD loop structure of S-Delta, accompanied by the reposition of the N-glycan on N149 (**Fig. 2A** and **Extended Data Fig. 6A**). The conformational changes in NTD loops and glycan shield provide an explanation for the escape from the NTD-specific nAbs^21^ and the markedly reduced efficacies of the convalescent and vaccine-elicited sera^25,26^.

In the same vein, the S-Gamma mutations that introduce the additional N-glycans on N20 and N188 introduce significant steric hindrance not only to the well-defined NTD supersite but also to an alternative epitope in turn blocking NTD-specific nAb binding (**Fig. 2C-F, Extended Data Fig. 6B** and **Extended Data Fig. 7**). The N20 glycosylation is indeed observed in the full-length S-Gamma based on cryo-EM data but the N188 glycosylation is not mentioned^36^. Additionally, this study only observed two distinct 1 RBD-up conformation for S-Gamma, while anther cryo-EM study also reported a single 1 RBD-up conformation^15^. In contrast, we observed an additional 2 RBD-up conformation (**Fig. 1D**). Of all the S variants investigated herein, S-Beta is the only one that exhibited a fully open 3 RBD-up conformation (**Fig. 1C**). Such a conformation resembles the structure of fully open S-D614G^38^, but it is not observed in previous studies^39,40^. The difference in the number of RBD-up domains may be attributed to the construct design and the experimental conditions. To address this issue, we further verified the authenticity of the 3 RBD-up conformation of S-Beta using the furin-cleavable form without the fm2P modification (**Fig. 1C** and **Extended Data Fig. 2**). Perhaps the most intriguing finding in terms of the S protein structure is the dimer of trimers of S-Kappa (**Fig. 1F** and **Fig. 3)**. While the biological implication of such a higher order assembly remains to be established, it is consistent with the unique E484Q mutation in the Kappa and other rare lineages.

Although the reported ACE2 binding affinities vary considerably depending on the construct designs of ACE2 (monomeric versus dimeric Fc-fusion^36^ or GFP-fusion used herein) and S proteins (full-length, trimeric ectodomain or monomeric RBD^36,37,39,40^, our self-consistent BLI analysis showed a strong correlation between the free energy derived from the K_d_ values and the size of the ACE2 binding interface area (**Fig. 4D**) based on the high-resolution cryo-EM structures of individual S variants (**Fig. 4B, 4C, and Extended Data Figs 8-12**). Furthermore, enhanced ACE2 binding can be rationalized by changes in the isoelectrostatic potential surface around the ACE2 binding site, which is highly modulated by the individual mutations (**Fig. 4E**).

NTD mutations in the Beta, Gamma, Delta and Kappa variant considerably attenuate various NTD-specific nAbs and completely abolish their neutralizing activities in many cases^15,17,20,21,27,36,37^. Therefore, nAbs that target the RBD remain a preferred route for success, and several RBD-specific nAbs are currently approved for emergency use to treat COVID-19 patients^12^. In this context, RBD-chAb-15, −25 and −45 that we investigated here are highly potent in neutralizing all four variants with the exception for RBD-chAb-25, which is rendered ineffective due to the N501Y mutation in S-Beta and S-Gamma (**Fig. 5**). Nevertheless, all three nAbs potently neutralize the most problematic Delta variant, as well as the Kappa variant. According to our previously reported high-resolution cryo-EM structures of RBD-chAb-15, −25 and −45 in complex with S variants^32,41^, RBD-chAb-45 binds to the site Ia/class 2 on RBD, which is accessible regardless of the RBD up/down conformation^8,12^, while RBD-chAb-15 and −25 bind to different regions within the epitope Ib/class 1 on RBD, which is only accessible in the RBD-up state^8,12^. The versatility of RBD-chAb-45 in recognizing different states of the RBD is manifested in the lowest IC_50_ values against all four pseudovirus variants (**Fig. 5C**). Furthermore, RBD-chAb-15 and −45 simultaneously and non-competitively bind to all three RBDs with a 3:3:3 stoichiometry (**Fig. 5B** and **Extended Data Fig. 13A**). Combined with our previous studies on the same set of nAbs against the original Wuhan strain^41^ and the Alpha variant^32^, our current study demonstrates the broad neutralization capacity and therapeutic potential of using a cocktail of these novel nAbs.

### Associated content

The atomic coordinates of the S variants in apo form and in complex with ACE2 are deposited in the Protein Data Bank (PDB) under the accession codes 7V76 to 7V77,7V8C (S-Beta), 7V78 to 7V7A (S-Gamma), 7V7N to 7V7V (S-Delta), 7V7D to 7V7J (S-Kappa), 7V7Z to 7V80 (S-Beta:ACE2), 7V81 to 7V84 (S-Gamma:ACE2), 7V88 to 7V8B (S-Delta:ACE2) and 7V85 to 7V87 (S-Kappa:ACE2). The cryo-EM maps are deposited in the Electron Microscopy Data Bank (EMDB) under successive codes from EMD-31760, EMD-31761, EMD-31797 to 31800 (S-Beta), EMD-31762 to 31764 (S-Gamma), EMD-31775 to 31783 (S-Delta), EMD-31767 to 31773 (S-Kappa), EMD-31784 to 31785 (S-Beta:ACE2), EMD-31786 to 31789 (S-Gamma:ACE2), EMD-31793 to 31796 (S-Delta:ACE2) and EMD-31790 to 31792 (S-Kappa:ACE2). The deposited EMDB codes of NSEM maps of the S variants in complex with RBD-chAb-15/45 cocktail are EMD-31817 (for S-Beta), EMD-31818 (for S-Gamma), EMD-31819 (for S-Delta) and EMD-31820 (S-Kappa), and in complex with RBD-chAb-25 are EMD-31821 (for S-Delta) and EMD-31822 (for S-Kappa). The LC-MS/MS raw datasets of S-Beta, S-Gamma, S-Delta and S-Kappa have been deposited to MassIVE site under the accession code MSV000088012.

## Supporting information

Extended Data Figs. 1-14, Extended Data Tables 1-14 and Extended Data Movies 1-4 Supplementary Information Table 1

## Funding sources

This research was supported by the intramural funding Academia Sinica for S.T.D.H., K.H.K, and H.C.W., an Academia Sinica Career Development Award (AS-CDA-109-L08) and an Infectious Disease Research Supporting Grant (AS-IDR-110-08) to S.T.D.H., and the research grants from the Ministry of Science and Technology, Taiwan (MOST-108-3114-Y-001-002 and MOST-108-2823-8-001-001 to H.C.W, and MOST 109-3114-Y-001-001 to H.C.W. and S.T.D.H.).

## Acknowledgement

We thank the Academia Sinica Cryo-EM Center (AS-CFII-108-110), Academia Sinica Biophysics Core Facility (AS-CFII108-111), Academia Sinica Common Mass Spectrometry Facilities for Proteomics and Protein Modification Analysis (AS-CFII-108-107) for data collection, and the Academia Sinica RNAi Core Facility for providing SARS-CoV-2 pseudotyped lentiviruses, all of which are funded by the Academia Sinica Core Facility and Innovative Instrument Project. The Academia Sinica Cryo-EM Center is also supported by the Taiwan Protein Project (AS-KPQ-109-TPP2). We thank the mammalian cell culture facility of the Institute of Biological Chemistry, Academia Sinica, for supporting protein production, the Academia Sinica Grid Computing for cryo-EM data processing, and Yun Mou at the Institute of Biomedical Sciences, Academia Sinica, for sharing the BLI instrument.

## Author contributions

S.T.D.H. conceived the project. T.J.Y. and P.Y.Y. prepared the S proteins and ACE2. H.C.W. provided the RBD-chAbs. T.J.Y. and Y.C.C. collected the cryo-EM data. T.J.Y. processed the cryo-EM data and determined the atomic structures with inputs from P.D.. T.J.Y. and P.Y.Y. collected and analyzed the NSEM and BLI data. K.H.L collected and analyzed pseudovirus data. N.E.C., Y.X.T. and Y.S.W. collected and analyzed the MS glycosylation data with inputs from Y.C.C. and K.H.K. to develop the analytical workflow. T.J.Y., P.Y.Y. and B.L. curated the nAb and clinical data. T.J.Y., P.Y.Y. and S.T.D.H. wrote the manuscript with inputs from all co-authors. K.H.K., H.C.W. and S.T.D.H. obtained funding.

## Competing interests

Related to this work, the Academia Sinica has filed a US patent application for the neutralizing antibodies on which H.-C.W. and S.-T.D.H. are named as inventors. The patent application is pending approval. The other authors declare no conflict of interest.

## Methods

### Expression and purification of S-Beta, Gamma, Delta, and Kappa variants

The codon-optimized DNA sequences corresponding to residues 1-1209 of S-Beta, Gamma, Delta, and Kappa variants were individually cloned into the mammalian expression vector pcDNA3.4-TOPO (Invitrogen, U. S. A.), which contains a foldon trimerization domain based on phage T4 fibritin followed by a c-Myc epitope and a hexa-repeat histidine tag as previously described^32,42^. All constructs contained the fm2P modification, which is defined as the tandem proline replacement (2P, ^986^KV^987^ → ^986^PP^987^) and the furin cleavage site mutation (fm, ^682^RRAR^685^ → ^682^GSAG^685^), for stabilizing the S protein in a prefusion state^7^. The plasmids of all S variants were transiently expressed in HEK293 Freestyle cells with polyethylenimine. Recombinant protein purification was carried out as previously described^32^. Briefly, the culture medium was incubated with HisPur Cobalt Resin (Thermo Fisher Scientific, U. S. A.) in binding buffer (50 mM Tris-HCl (pH 7.6), 300 mM NaCl, 10 mM imidazole, and 0.02% NaN_3_) at 4ºC overnight. The resin was thoroughly washed with wash buffer (50 mM Tris-HCl (pH 7.6), 300 mM NaCl, and 10 mM imidazole). The recombinant S proteins were eluted by elution buffer (50 mM Tris-HCl (pH 7.6), 150 mM NaCl, and 150 mM imidazole). The eluted S proteins were further purified by size-exclusion chromatography (SEC) using a Superose 6 increase 10/300 GL column (GE Healthcare, U. S. A.) in TBS buffer (50 mM Tris-HCl (pH 7.6), 150 mM NaCl, and 0.02 % NaN_3_). The protein concentrations were determined using the UV absorbance at 280 nm using a UV-Vis spectrometer (Nano-photometer N60, IMPLEN, Germany).

### Expression and purification of sfGFP-ACE2

The production of recombinant sfGFP-ACE2 (hereafter ACE2) was described previously^32^. Briefly, transient expression of recombinant ACE2 was achieved in Expi293 cells. The culture medium containing the secreted ACE2 was subsequently loaded onto an ion-exchange column (HiTrap 5 ml Q FF anion exchange chromatography column; GE Healthcare, U. S. A.), and subsequently eluted by a linear salt gradient ranging between 10 and 1000 mM NaCl. A final polishing step by a Superdex 75 16/600 GL size-exclusion column (GE Healthcare, U. S. A.) was carried out in TBS buffer. The protein concentrations were determined by using the UV absorbance at 280 nm using a UV-Vis spectrometer (Nano-photometer N60, IMPLEN, Germany).

### Biolayer interferometry (BLI)

ACE2 was biotinylated and immobilized onto the High Precision Streptavidin (SAX) biosensors (Sartorius, Germany) in assay buffer (50 mM Tris-HCl (pH 7.6), 150 mM NaCl, 0.02% NaN_3_, and 0.1% BSA). S-Beta, S-Gamma, S-Delta, and S-Kappa were serially diluted to 100, 50, 25, 12.5, 6.25, and 3.125 nM in assay buffer for independent binding using an OctetRED 96 biolayer interferometer (ForteBio, U. S. A.). All measurements were conducted at 25 ºC. The association and dissociation periods were set to 600 and 900 s, respectively. The independent BLI sensorgrams with different S protein concentrations were baseline-corrected using double reference as previously described^32^. The double-reference subtracted data were globally fit to a 1:1 binding model using the built-in Data Analysis v10.0 software (ForteBio, U. S. A.). For RBD-chAb neutralization analysis, RBD-chAb-15, −25, and −45 were individually pre-incubated with an equal molar ratio of S-Beta, S-Gamma, S-Delta, or S-Kappa at a final concentration of 50 nM at room temperature for 1 hr. The measurements were carried out by using the same ACE2-immobilized biosensors with 400 s of association and 150 s of dissociation. The processed data were exported and replotted by using Prism 9 (GraphPad, U. S. A.).

### Negative staining electron microscopy (NSEM) analysis

For all NSEM samples, four microliters of protein solution at a concentration of 50 μg/mL were used to prepare NSEM grids. The samples included S-Delta and S-Kappa in complex with RBD-chAb-25, and all four S variants (S-Beta, S-Gamma, S-Delta and S-Kappa) in complex with equal amounts of RBD-chAb-15 and −45. The carbon-coated grids were glow-discharged at 25 mA for 30 s. After staining with 0.2% uranyl formate (UF) and blotting by filter paper, the grids were then air-dried for one day. All images were collected by using a FEI Tecnai G2-F20 electron microscope at 200 keV (FEI, the Netherlands) with a magnification of 50,000x. The resulting pixel size was 1.732 Å. Data processing was accomplished by using cryoSPARC v2.14. After patch-CTF estimation, particle images were picked by using the function of “blob picker”, and extracted with a box size of 256 pixels. Particle clean-up was achieved by iterative rounds of 2D classification. Selected particles from the best 2D classes were further used for ab-initio 3D reconstruction and homogeneous refinement. The resulting NSEM maps were visualized and analyzed by UCSF-ChimeraX^43^.

### Cryo-EM sample preparation and Data collection

For all cryo-EM grids, three microliters of protein solution at a concentration of 1.5 mg/mL were applied onto 300-mesh Quantifoil R1.2/1.3 holey carbon grids. The grids were glow-discharged at 20 mA for 30 s. After a 30-s waiting time, the grids were blotted for 2.5 s at 4 ºC with 100% humidity, and vitrified using a Vitrobot Mark IV (ThermoFisher Scientific, U. S. A.). The samples included apo S variants, and S variants in complex with ACE2 (each S variant was pre-incubated with ACE2 at a molar ratio of 1:1.3 at room temperature for 1 hr prior to vitrification).

Data acquisition was performed on a 300 keV Titan Krios microscope equipped with a Gatan K3 direct electron detector (Gatan, U. S. A.) in a super-resolution mode using EPU v2.10 software (ThermoFisher Scientific, U. S. A.). Movies were collected with a defocus range between −0.8 and −2.6 µm with a magnification of 81000 x, resulting in a pixel size of 0.55 Å. A total dose of 45-50 e^-^/Å^2^ was distributed over 50 frames with an exposure time of 1.8 s. The dataset was collected with an energy filter (slit width: 15 eV), and the dose rate was adjusted to 8-9 e^-^/pix/s.

### Image processing and 3D reconstruction

All 2x binned super-resolution movie files were analyzed by Relion-3.0^44^ with dose-weighting and 5×5 patch-based alignment using MotionCor2 (v1.2.6)^45^. All motion-corrected micrographs were transferred to cryoSPARC v2.14 ^46^ for further processing. Contrast transfer function (CTF) estimation was achieved by patch-based CTF. The micrographs that have the “CTF_fit_to_Res” parameters between 2.5 and 4 Å were selected for subsequent particle picking. The picked particles were Fourier-cropped to a box size of 192 pixels after the particle extraction with the 384-pixels box size and then applied to iterative rounds of 2D classification for filtering junk particles.

In general, particle images were picked and classified by ab-initio reconstruction with C1 symmetry, followed by heterogeneous refinement to generate five distinct classes. Next, the particles from the best 3D classes were applied to non-uniform (NU) refinement to generate an initial model. The initial 3D model with a refined mask from the NU refinement was used for three-dimensional variability analysis (3DVA). Five clusters generated from the 3DVA as the templates were used for further heterogeneous refinement. Particles corresponding to three of the five clusters were re-extracted, un-binned, and non-uniformly refined with local CTF refinement to yield the final cryo-EM maps (**Extended Data Figs. 2-3**).

For S-Delta and S-Kappa. For S-Delta, in addition to the above image processing procedures, another round of 3DVA and heterogeneous refinement were necessary to improve the definitions of the individual RBD maps. This resulted in a total of nine cryo-EM maps for S-Delta (**Extended Data Fig. 4**). For S-Kappa, 2D classification and initial heterogeneous refinement revealed two distinct classes, with one being the canonical S trimer with the one RBD-up conformation and the other being an unprecedented dimer of S trimers. The two sets of particles were analyzed separately. For the canonical S-Kappa trimer, two rounds of 3DVA and heterogeneous refinement were performed as did for S-Delta. These steps teased out a set of particles that correspond to the dimer of S trimers so they were joined with another particle pool to determine the atomic structures of the dimer of S trimers. All particles from different 3D classes were further re-extracted with a box size of 384 pixels and applied to NU refinement to generate seven cryo-EM maps for S-Kappa, including for canonical trimeric S with different RBD conformations and three for dimer of S trimers (**Extended Data Fig. 5**).

A similar workflow was applied to ACE2-bound S variants. For S-Beta: ACE2, a 3.2 Å cryo-EM map of S-Beta engaged with ACE2 in a 3:3 stoichiometry (3 ACE2-bound) was obtained (**Extended Data Fig. 8**). For S-Gamma, S-Delta and S-Kappa:ACE2, a 2 ACE2-bound population was separated from the dominant 3 ACE2-bound population to generate separate sets of cryo-EM maps (**Extended Data Figs. 9-11**). To improve the resolution of the binding interface between RBD and ACE2 for all S variants, another round of local refinement with a focused mask covering the interface was performed. The focus masks were created by UCSF-ChimeraX^43^. This yielded a separate set of cryo-EM maps of focus-refined ACE2 in complex with RBD with nominal resolutions ranged between 3.0 and 3.0 Å (**Extended Data Figs. 8-11**).

### Model building and refinement

Initial models of S-Beta, S-Gamma, S-Delta, and S-Kappa was generated based on the previously reported structure of S-D614G (PDB ID: 7EB3) aided by Swiss-Model for in silico mutagenesis^33,47^. The atomic coordinates were divided into individual domains and manually fitted into the cryoEM map in UCSF-ChimeraX^43^, and manually optimized by Coot^48^. The same procedure was applied to S variants in complex with ACE2. The atomic structure of ACE2 in complex with S-Alpha (PDB ID: 7EDJ) was used as the initial model^32^. After iterative refinements, the models were further processed by real-space refinement in Phenix^49^ to attain convergent models. N-linked glycans based on the N-glycosylation sequon (<u>N</u>-X-S/T) were added onto asparagine side-chains by using the extension module “Glyco” within Coot^48^. The final models were assessed by a combinative tool of comprehensive cryo-EM validation in Phenix^49,50^. Structural visualization and rendering of structural representations were accomplished by using UCSF-ChimeraX and Pymol 2.4.1 (Schrodinger Inc. U. S. A.).

### Structure-based statistics of N-terminal domain (NTD) epitope binding frequency

The statistics of residue-specific epitope usage of NTD-specific nAbs was done by using the ‘Protein interfaces, surfaces and assemblies’ service PISA at the European Bioinformatics Institute (http://www.ebi.ac.uk/pdbe/prot_int/pistart.html)^51^. The list of cryo-EM and crystal structures of nAb-bound spike or truncated NTD was manually curated by searching the EMDB/PDB entries deposited in the Protein Data Bank and their corresponding publications as tabulated in *Extended Table 7*. The nAb contacting residues within NTD were defined by the function “Interfaces” of PISA and summarized in *Supplementary information Table 1*. The cumulated epitope counts were mapped onto the homology model of NTD generated by Swiss-model^47^, and rendered by using PyMol 2.4.1 (Schrodinger Inc. U. S. A.).

### Pseudovirus neutralization assay

The pseudovirus neutralization assays were performed using 293T cells overexpressing ACE2 as described previously^32^. The pseudoviruses of SARS-CoV-2 variants were prepared the by RNAi Core of Academia Sinica. Four-fold serially dilution of chAbs were premixed with 1000 TU/well Beta, Gamma, Delta, and Kappa strains of SARS-CoV-2 pseudovirus. The mixture was incubated for 1 h at 37°C and then added to pre-seeded 293T-ACE2 cells at 100 μl/well for 24 h at 37°C. The medium was removed and refilled with 100 μl/well DMEM for additional 48-h incubation. Next, 100 μl ONE-Glo™ luciferase reagent (Promega) was added to each well for 3-min incubation at 37°C. The luciferase activity was determined using a microplate spectrophotometer (Molecular Devices). The inhibition rate was calculated by comparing the luminescence value to the negative and positive control wells. IC_50_ was determined by a four-parameter logistic regression using GraphPad Prism 9 (GraphPad, U. S. A.).

### Glycosylation analysis by mass spectrometry

The in-solution proteolytic digestion, LC-MS/MS analysis, glycopeptide identification and quantification of S-Beta, S-Gamma, S-Delta, S-Kappa were carried out by using the methods described previously^52^, without searching for O-glycopeptide. N-glycopeptide identification was achieved by Byonic and the quantitation was achieved by Byos (Protein Metrics Inc., USA). The built-in N-glycan library of “132 human” for N-glycopeptide identification was used with no further modifications.

### Atomic model building of fully glycosylated NTD

To define the representative glycoform for individual N-glycosylation sites on the NTD of S-Gamma, the most abundant glycoforms derived from the aforementioned mass spectrometry analyses were used. In cases where the glycoforms could not be defined experimentally, i.e., N17 and N149 for S-Gamma, the glycoforms used in a previously reported atomistic molecular dynamics simulation study^53^ were used. In the case of N20, which is unique to S-Gamma, the same glycoform as that of N17 was used. The atomic coordinates of the individual N-glycans were manually built by using Coot^48^ following the same procedure as described previously^42^.

## References

1 Outbreak.info. a standardized, open-source database of COVID-19 resources and epidemiology data, https://outbreak.info (2021).

2 World Health Organization. Tracking SARS-CoV-2 variants, https://www.who.int/en/activities/tracking-SARS-CoV-2-variants/ (2021).

3 Centers for Disease Control and Prevention. SARS-CoV-2 Variant Classifications and Definitions, https://www.cdc.gov/coronavirus/2019-ncov/variants/variant-info.html (2021).

4 Faria, N. R. et al. Genomics and epidemiology of the P.1 SARS-CoV-2 lineage in Manaus, Brazil. Science 372, 815–821, doi:10.1126/science.abh2644 (2021).

5 Funk, T. et al. Characteristics of SARS-CoV-2 variants of concern B.1.1.7, B.1.351 or P.1: data from seven EU/EEA countries, weeks 38/2020 to 10/2021. Euro Surveill 26, doi:10.2807/1560-7917.ES.2021.26.16.2100348 (2021).

6 Walls, A. C. et al. Structure, Function, and Antigenicity of the SARS-CoV-2 Spike Glycoprotein. Cell 181, 281–292 e286, doi:10.1016/j.cell.2020.02.058 (2020).

7 Wrapp, D. et al. Cryo-EM structure of the 2019-nCoV spike in the prefusion conformation. Science 367, 1260–1263, doi:10.1126/science.abb2507 (2020).

8 Barnes, C. O. et al. SARS-CoV-2 neutralizing antibody structures inform therapeutic strategies. Nature 588, 682–687, doi:10.1038/s41586-020-2852-1 (2020).

9 Cao, Y. et al. Potent Neutralizing Antibodies against SARS-CoV-2 Identified by High-Throughput Single-Cell Sequencing of Convalescent Patients’ B Cells. Cell 182, 73–84 e16, doi:10.1016/j.cell.2020.05.025 (2020).

10 Piccoli, L. et al. Mapping Neutralizing and Immunodominant Sites on the SARS-CoV-2 Spike Receptor-Binding Domain by Structure-Guided High-Resolution Serology. Cell 183, 1024–1042 e1021, doi:10.1016/j.cell.2020.09.037 (2020).

11 Liu, L. et al. Potent neutralizing antibodies against multiple epitopes on SARS-CoV-2 spike. Nature 584, 450–456, doi:10.1038/s41586-020-2571-7 (2020).

12 Corti, D., Purcell, L. A., Snell, G. & Veesler, D. Tackling COVID-19 with neutralizing monoclonal antibodies. Cell 184, 4593–4595, doi:10.1016/j.cell.2021.07.027 (2021).

13 U.S FOOD & DRUG. Coronavirus (COVID-19) Update: FDA Revokes Emergency Use Authorization for Monoclonal Antibody Bamlanivimab, https://www.fda.gov/news-events/press-announcements/coronavirus-covid-19-update-fda-revokes-emergency-use-authorization-monoclonal-antibody-bamlanivimab (2021).

14 Watanabe, Y., Allen, J. D., Wrapp, D., McLellan, J. S. & Crispin, M. Site-specific glycan analysis of the SARS-CoV-2 spike. Science 369, 330–333, doi:10.1126/science.abb9983 (2020).

15 Wang, P. et al. Increased resistance of SARS-CoV-2 variant P.1 to antibody neutralization. Cell Host Microbe 29, 747–751 e744, doi:10.1016/j.chom.2021.04.007 (2021).

16 Chi, X. et al. A neutralizing human antibody binds to the N-terminal domain of the Spike protein of SARS-CoV-2. Science 369, 650–655, doi:10.1126/science.abc6952 (2020).

17 McCallum, M. et al. N-terminal domain antigenic mapping reveals a site of vulnerability for SARS-CoV-2. Cell 184, 2332–2347 e2316, doi:10.1016/j.cell.2021.03.028 (2021).

18 Rosa, A. et al. SARS-CoV-2 can recruit a heme metabolite to evade antibody immunity. Sci Adv 7, doi:10.1126/sciadv.abg7607 (2021).

19 Cerutti, G. et al. Potent SARS-CoV-2 neutralizing antibodies directed against spike N-terminal domain target a single supersite. Cell Host Microbe 29, 819–833 e817, doi:10.1016/j.chom.2021.03.005 (2021).

20 Wang, P. et al. Antibody resistance of SARS-CoV-2 variants B.1.351 and B.1.1.7. Nature 593, 130–135, doi:10.1038/s41586-021-03398-2 (2021).

21 Liu, Y. et al. The SARS-CoV-2 Delta variant is poised to acquire complete resistance to wild-type spike vaccines. biorxiv, doi:10.1101/2021.08.22.457114 (2021).

22 Caniels, T. G. et al. Emerging SARS-CoV-2 variants of concern evade humoral immune responses from infection and vaccination. medRxiv, doi:10.1101/2021.05.26.21257441 (2021).

23 Tada, T. et al. Convalescent-Phase Sera and Vaccine-Elicited Antibodies Largely Maintain Neutralizing Titer against Global SARS-CoV-2 Variant Spikes. mBio 12, e0069621, doi:10.1128/mBio.00696-21 (2021).

24 Zhou, H. et al. B.1.526 SARS-CoV-2 variants identified in New York City are neutralized by vaccine-elicited and therapeutic monoclonal antibodies. mBio, doi:10.1101/2021.03.24.436620 (2021).

25 Choi, A. et al. Serum Neutralizing Activity of mRNA-1273 against SARS-CoV-2 Variants. N Engl J Med 384, 1468–1470, doi:10.1056/NEJMc2102179 (2021).

26 Carreño, J. M. et al. Reduced neutralizing activity of post-SARS-CoV-2 vaccination serum against variants B.1.617.2, B.1.351, B.1.1.7+E484K and a sub-variant of C.37. medRxiv, doi:10.1101/2021.07.21.21260961 (2021).

27 Mlcochova, P. et al. SARS-CoV-2 B.1.617.2 Delta variant replication, sensitivity to neutralising antibodies and vaccine breakthrough. bioRxiv, doi:10.1101/2021.05.08.443253 (2021).

28 Liu, J. et al. BNT162b2-elicited neutralization of B.1.617 and other SARS-CoV-2 variants. Nature 596, 273–275, doi:10.1038/s41586-021-03693-y (2021).

29 Planas, D. et al. Reduced sensitivity of infectious SARS-CoV-2 variant B.1.617.2 to monoclonal antibodies and sera from convalescent and vaccinated individuals. Nature 596, 276–280, doi:10.1101/2021.05.26.445838 (2021).

30 Liu, C. et al. Reduced neutralization of SARS-CoV-2 B.1.617 by vaccine and convalescent serum. Cell 184, 4220–4236 e4213, doi:10.1016/j.cell.2021.06.020 (2021).

31 Lopez Bernal, J. et al. Effectiveness of Covid-19 Vaccines against the B.1.617.2 (Delta) Variant. N Engl J Med 385, 585–594, doi:10.1056/NEJMoa2108891 (2021).

32 Yang, T. J. et al. Effect of SARS-CoV-2 B.1.1.7 mutations on spike protein structure and function. Nat. Struct. Mol. Biol., doi:10.1038/s41594-021-00652-z (2021).

33 Yang, T.-J. et al. COVID-19 dominant D614G mutation in the SARS-CoV-2 spike protein desensitizes its temperature-dependent denaturation. bioRxiv, doi:10.1101/2021.03.28.437426 (2021).

34 Corti, D., Purcell, L. A., Snell, G. & Veesler, D. Tackling COVID-19 with neutralizing monoclonal antibodies. Cell 184, 3086–3108, doi:10.1016/j.cell.2021.05.005 (2021).

35 Harvey, W. T. et al. SARS-CoV-2 variants, spike mutations and immune escape. Nat Rev Microbiol 19, 409–424, doi:10.1038/s41579-021-00573-0 (2021).

36 Zhang, J. et al. Membrane fusion and immune evasion by the spike protein of SARS-CoV-2 Delta variant. bioRxiv, doi:10.1101/2021.08.17.456689 (2021).

37 McCallum, M. et al. Molecular basis of immune evasion by the delta and kappa SARS-CoV-2 variants. bioRxiv, doi:10.1101/2021.08.11.455956 (2021).

38 Yurkovetskiy, L. et al. Structural and Functional Analysis of the D614G SARS-CoV-2 Spike Protein Variant. Cell 183, 739–751 e738, doi:10.1016/j.cell.2020.09.032 (2020).

39 Gobeil, S. M. et al. Effect of natural mutations of SARS-CoV-2 on spike structure, conformation, and antigenicity. Science 373, doi:10.1126/science.abi6226 (2021).

40 Cai, Y. et al. Structural basis for enhanced infectivity and immune evasion of SARS-CoV-2 variants. Science 373, 642–648, doi:10.1126/science.abi9745 (2021).

41 Su, S. C. et al. Structure-guided Antibody Cocktail for Prevention and Treatment of COVID-19. Plos Pathog (Accepted).

42 Yang, T. J. et al. Cryo-EM analysis of a feline coronavirus spike protein reveals a unique structure and camouflaging glycans. Proc Natl Acad Sci U S A 117, 1438–1446, doi:10.1073/pnas.1908898117 (2020).

43 Goddard, T. D. et al. UCSF ChimeraX: Meeting modern challenges in visualization and analysis. Protein Sci 27, 14–25, doi:10.1002/pro.3235 (2018).

44 Zivanov, J. et al. New tools for automated high-resolution cryo-EM structure determination in RELION-3. Elife 7, doi:10.7554/eLife.42166 (2018).

45 Zheng, S. Q. et al. MotionCor2: anisotropic correction of beam-induced motion for improved cryo-electron microscopy. Nat Methods 14, 331–332, doi:10.1038/nmeth.4193 (2017).

46 Punjani, A., Rubinstein, J. L., Fleet, D. J. & Brubaker, M. A. cryoSPARC: algorithms for rapid unsupervised cryo-EM structure determination. Nature methods 14, 290–296, doi:10.1038/nmeth.4169 (2017).

47 Waterhouse, A. et al. SWISS-MODEL: homology modelling of protein structures and complexes. Nucleic Acids Res 46, W296–W303, doi:10.1093/nar/gky427 (2018).

48 Emsley, P., Lohkamp, B., Scott, W. G. & Cowtan, K. Features and development of Coot. Acta Crystallogr D Biol Crystallogr 66, 486–501, doi:10.1107/S0907444910007493 (2010).

49 Adams, P. D. et al. PHENIX: a comprehensive Python-based system for macromolecular structure solution. Acta crystallographica. Section D, Biological crystallography 66, 213–221, doi:10.1107/S0907444909052925 (2010).

50 Chen, V. B. et al. MolProbity: all-atom structure validation for macromolecular crystallography. Acta Crystallogr D Biol Crystallogr 66, 12–21, doi:10.1107/S0907444909042073 (2010).

51 Krissinel, E. & Henrick, K. Inference of macromolecular assemblies from crystalline state. J Mol Biol 372, 774–797, doi:10.1016/j.jmb.2007.05.022 (2007).

52 Kuo, C.-W. et al. Distinct shifts in site-specific glycosylation pattern of SARS-CoV-2 spike proteins associated with arising mutations in the D614G and Alpha variants. bioRxiv, doi:10.1101/2021.07.21.453140 (2021).

53 Woo, H. et al. Developing a Fully Glycosylated Full-Length SARS-CoV-2 Spike Protein Model in a Viral Membrane. J Phys Chem B 124, 7128–7137, doi:10.1021/acs.jpcb.0c04553 (2020).

