## Supporting information for "Structure-activity relationships of B.1.617 and other SARS-CoV-2 spike variants"

**This PDF file includes:**

Extended Data Figs. 1-14, Extended Tables 1-14, Extended Movies 1-4 and references
Captions for Movies 1-4

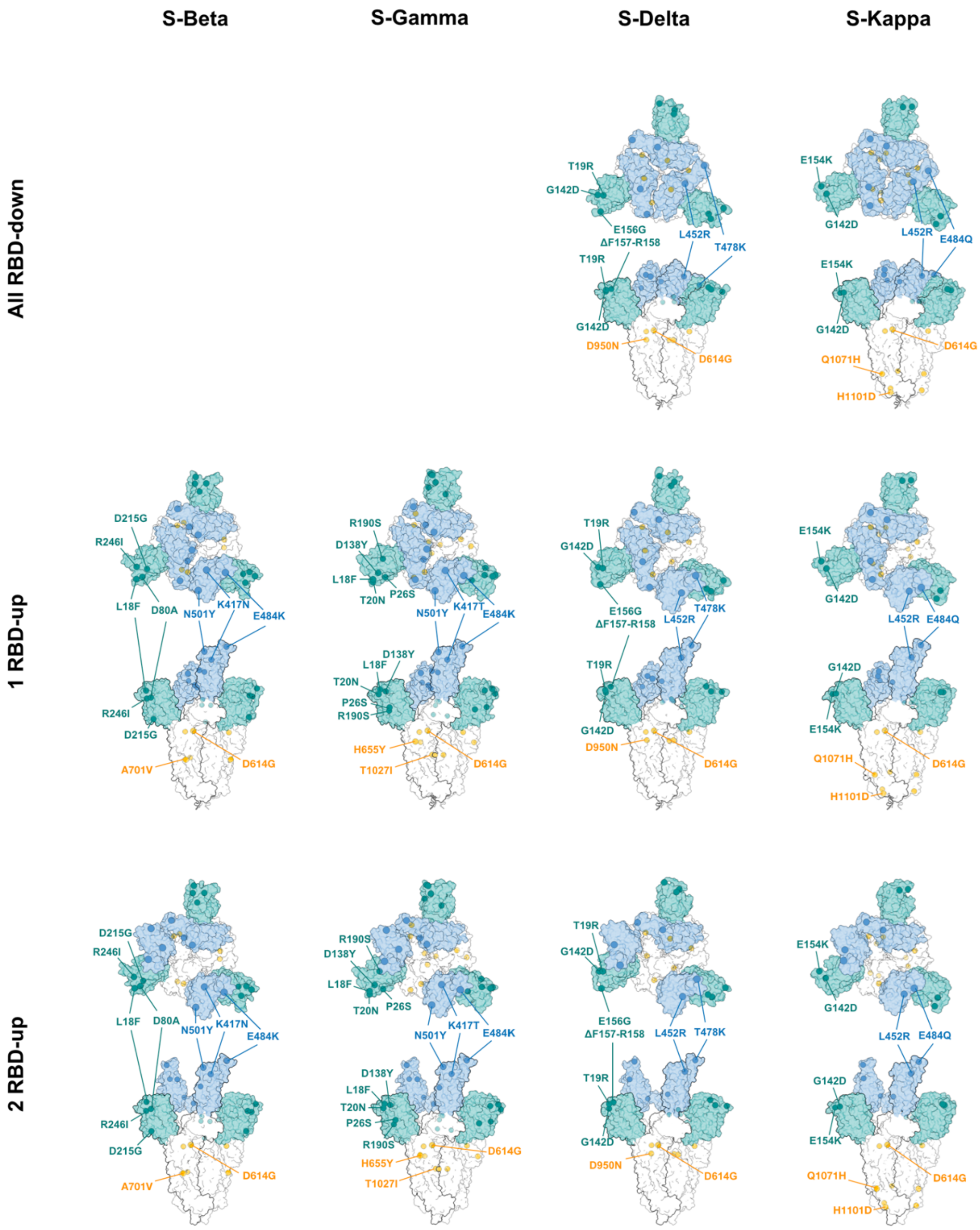

**Extended Data Fig. 1. Spatial distributions of variant-specific mutations of S-Beta, Gamma, Delta and Kappa**.

Orthogonal views of the representative cryo-EM structures in surface representation are shown for the distinct RBD states of individual S variants. NTD and RBD are colored accordingly to the coloring scheme in **Fig. 1A**. The backbone Cα atoms of the mutation sites are shown in spheres, with their identities for one of the RBD-up protomers, except for all-RBD down. The P681R mutation is not shown in S-Delta and S-Kappa due to the lack for resolved cryo-EM densities corresponding to the residues 674-688.

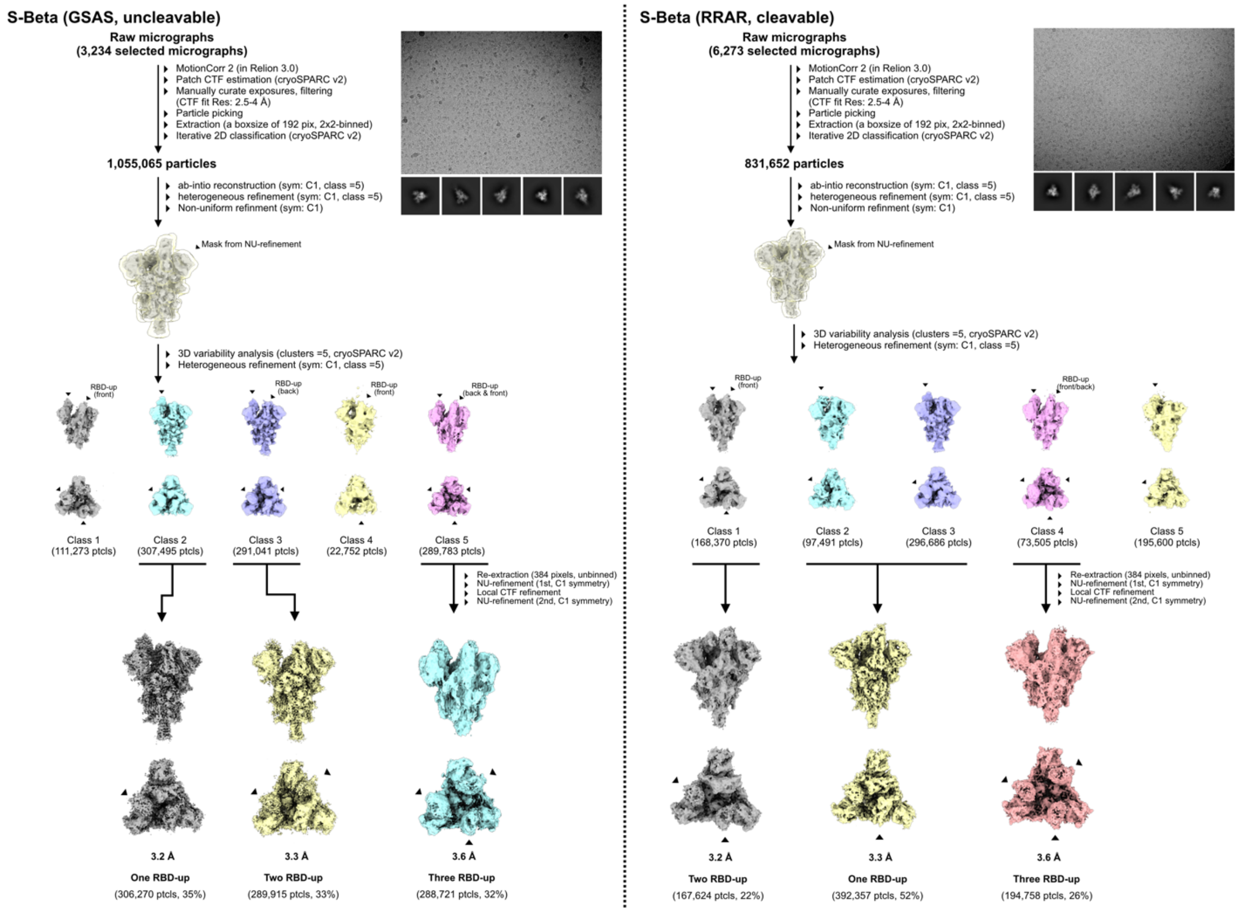

**Extended Data Fig. 2. Workflow for cryo-EM data processing of uncleavable and cleavable apo S-Beta.**

The motion-corrected micrographs were selected by the criteria of "CTF fit Res: 2.5-4 Å", and used for particles selection, and iterative rounds of 2D averaging classification. Particles from the best 2D classes were applied for initial 3D classification and non-uniform (NU) refinement. The resulting initial EM map from the NU refinement was used for 3DVA with a refined mask generated from the NU refinement, followed by another round of heterogeneous refinement with five clusters used as the templates. Particles from the 3D classes were re-extracted, re-grouped and subsequently applied to NU-refinement with the function of local CTF refinement to yield the final cryo-EM maps with the resolutions of 3.2-3.6 Å. The percentages of individual substates are indicated underneath the corresponding cryo-EM maps.

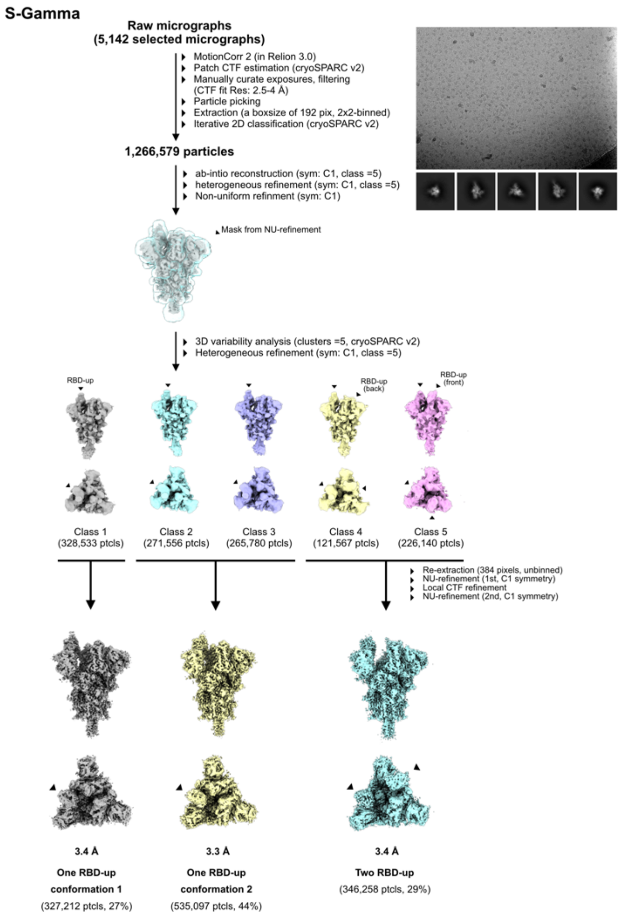

**Extended Data Fig. 3. Workflow for cryo-EM data processing of apo S-Gamma.**

The same selection criteria of micrographs and processing procedures described in *Extended Data Fig. 2* was used to generate five 3D classes after 3DVA. Particles from the 3D classes were re-extracted, re-grouped and subsequently applied to NU-refinement with the function of local CTF refinement to yield the final cryo-EM maps, including two 1 RBD-up and one 2 RBD-up maps with the resolutions of maps with 3.4, 3.3 and 3.4 Å, respectively.

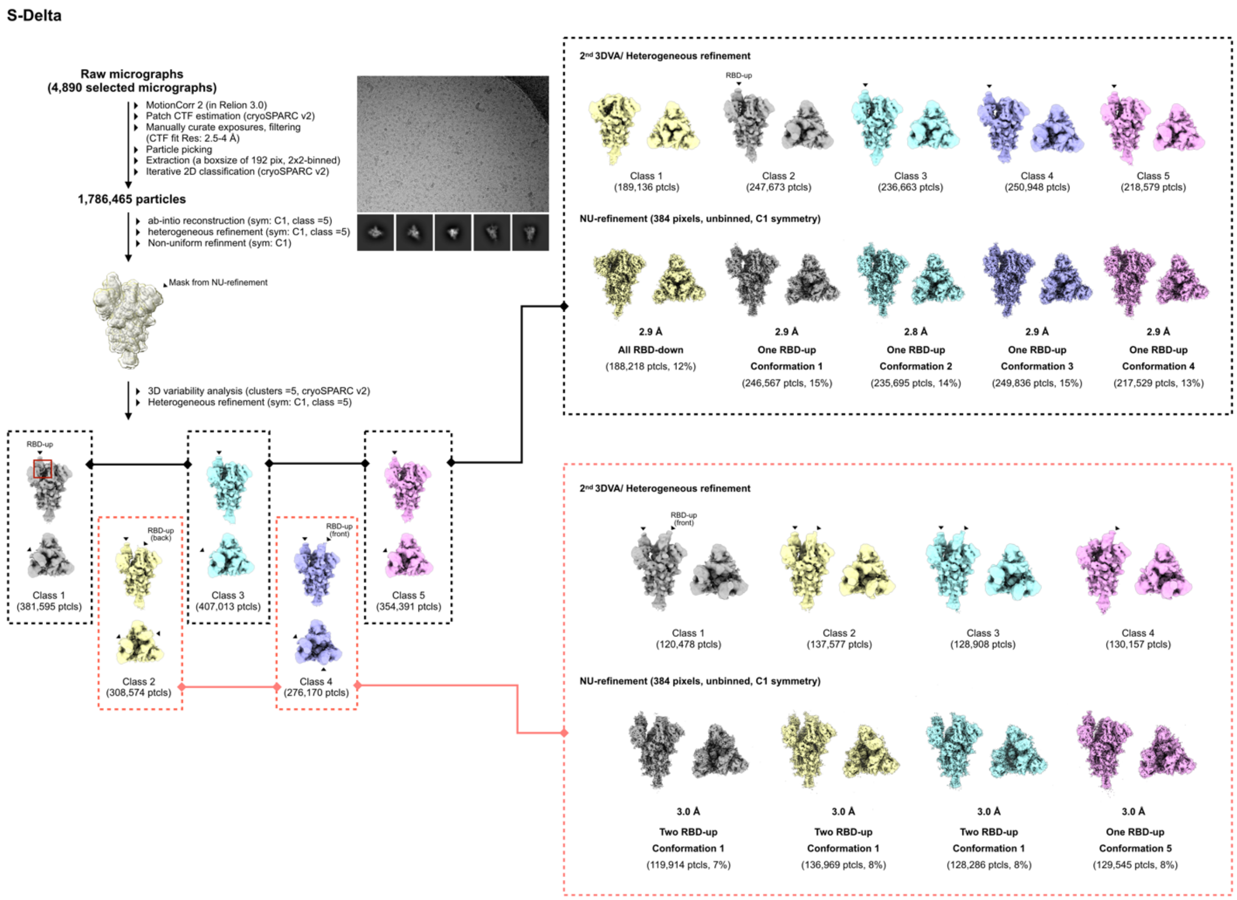

**Extended Data Fig. 4. Workflow for cryo-EM data processing of apo S-Delta.**

The same selection criteria of micrographs and processing procedures as described in *Extended Data Fig. 2* was used to generate five initial 3D classes. To improve the resolution of RBD region, another round of 3DVA followed by further heterogeneous refinement for each RBD-up conformation was carried out. Particles corresponding to different 3D classes were re-extracted and refined by NU-refinement, resulting in nine final cryo-EM maps, including one all-RBD, five 1 RBD-up, and three 2 RBD-up conformations.

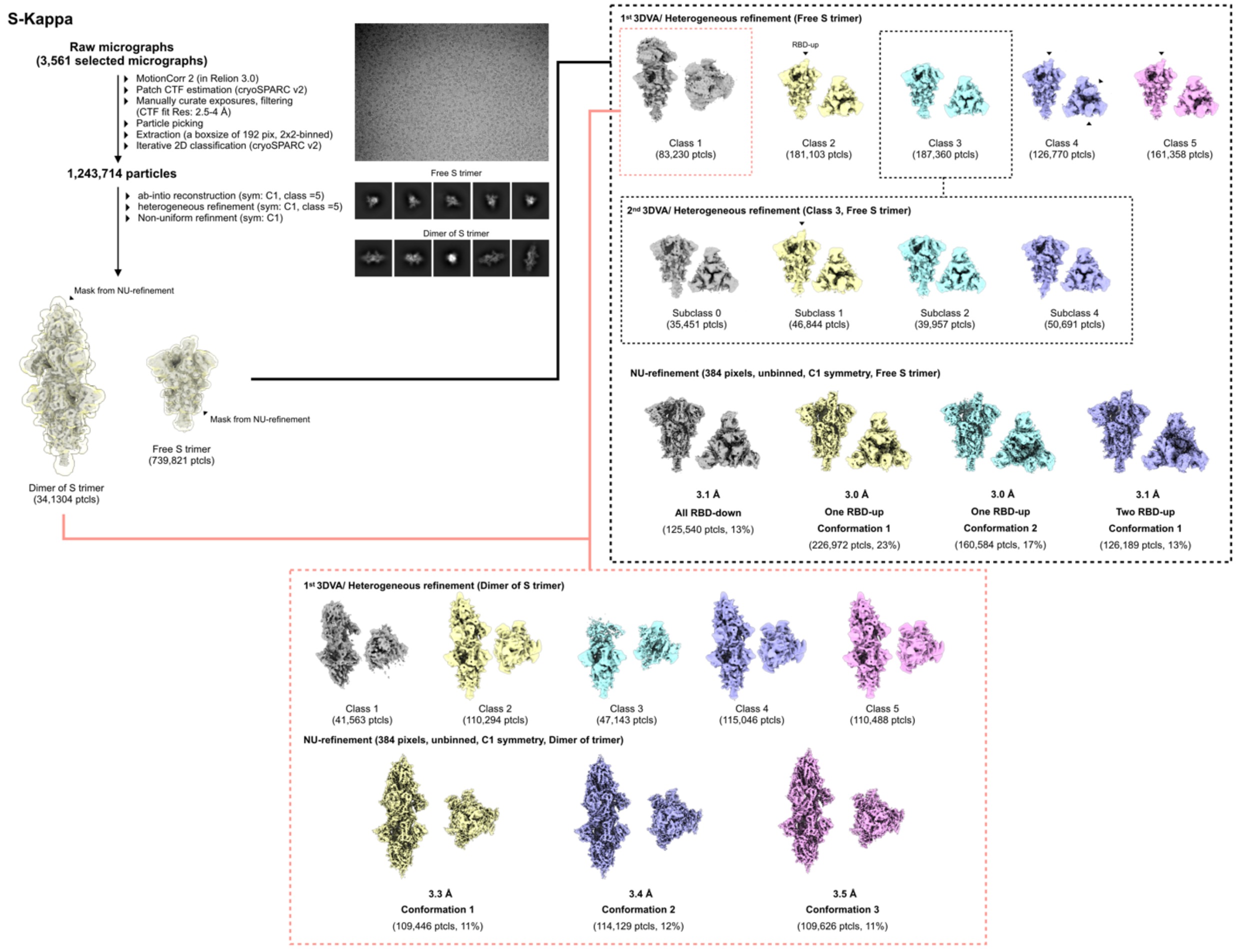

**Extended Data Fig. 5. Workflow for cryo-EM data processing of apo S-Kappa variant.**

The 2D classification and initial 3D refinement demonstrated the structures of the S-Kappa variant includes a class showing the canonical free trimer and another representing an unprecedented dimer of S-Kappa trimer. Two independent classes were further processed by 3DVA and subsequent heterogeneous refinement. The second round of 3DVA for particles of free S trimer was carried out to improve the 3D classification and resolution of the RBD region. All particles were re-extracted and refined by NU-refinement without symmetry (C1 symmetry) to yield seven cryo-EM structures, including one all RBD-down, two 1 RBD-up and one 2 RBD-up conformation, as well as three conformations of the dimer of S-Kappa trimers.

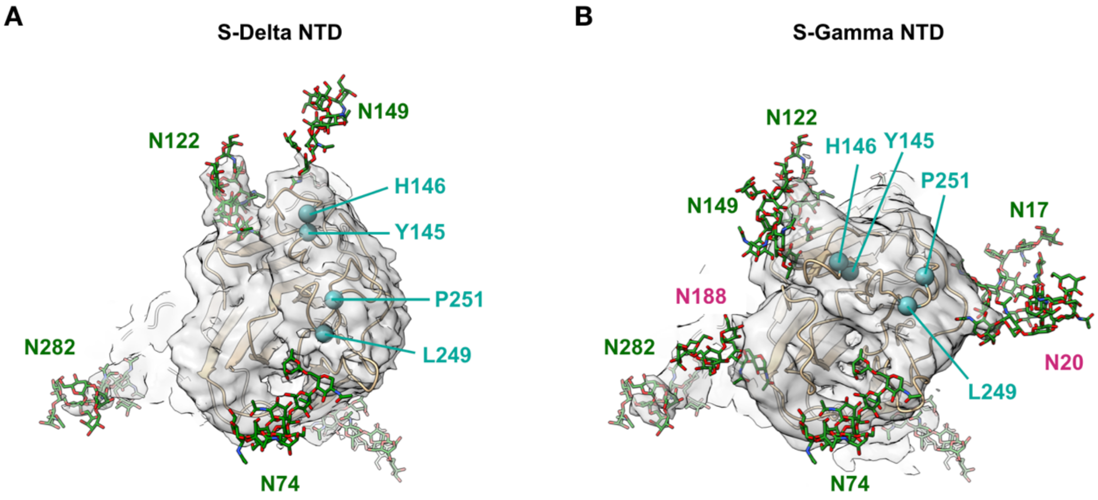

**Extended Data Fig. 6. Structural rearrangement of the NTD of S-Gamma and S-Delta.**

The Cartoon representation of the NTD of (A) S-Delta and (B) S-Gamma are overlaid by the corresponding cryo-EM maps (EMDB ID: EMD-P49J41 and EMD-P4626, respectively). The atomic models of the N-glycans are shown in sticks with carbon, nitrogen and oxygen atoms colored in green, blue and red, respectively. The positions of the Cα atoms of the most antigenic residues (supersites), i.e., Y145, H146, L249 and P251, are indicated by blue spheres. For S-Delta, the sequence follows the convention of S-WT despite the Δ157-158 deletions. For N122 and N282, most of the N-glycan models can be accounted for by the clear protrusions of EM maps extending from the respective asparagine side chains. For N17 and, N20 and N149, evidence of N-glycosylation can still be found as indicated by the smaller but significant protrusions extending from the respective asparagine side chains.

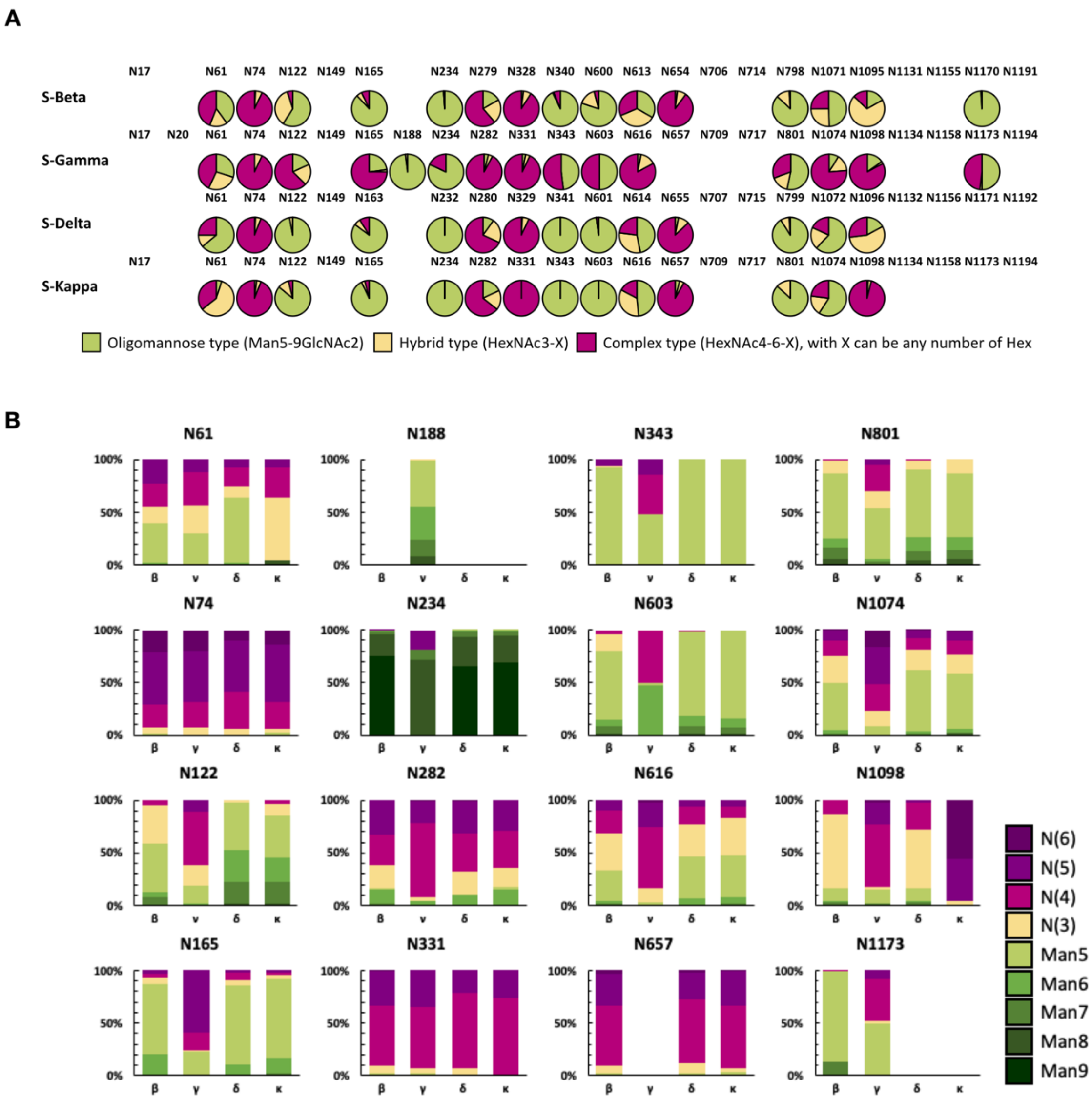

**Extended Data Fig. 7. Overall site-specific N-glycosylation patterns of S-Beta, S-Gamma, S-Delta and S-Kappa.**

(A) The pie charts summarize the quantification of the composition of N-glycosylation corresponding to the specific sites within each S variant. (B) Comparison of the full glycosylation heterogeneity of S-Beta, S-Gamma, S-Delta and S-Kappa. The bar charts showing the relative quantities of the high-mannose type N-glycan series and complex-type N-glycan series in each N-glycosylation site. The coloring scheme for different N-glycan types is indicated on the right, which follows the definition described previously [1].

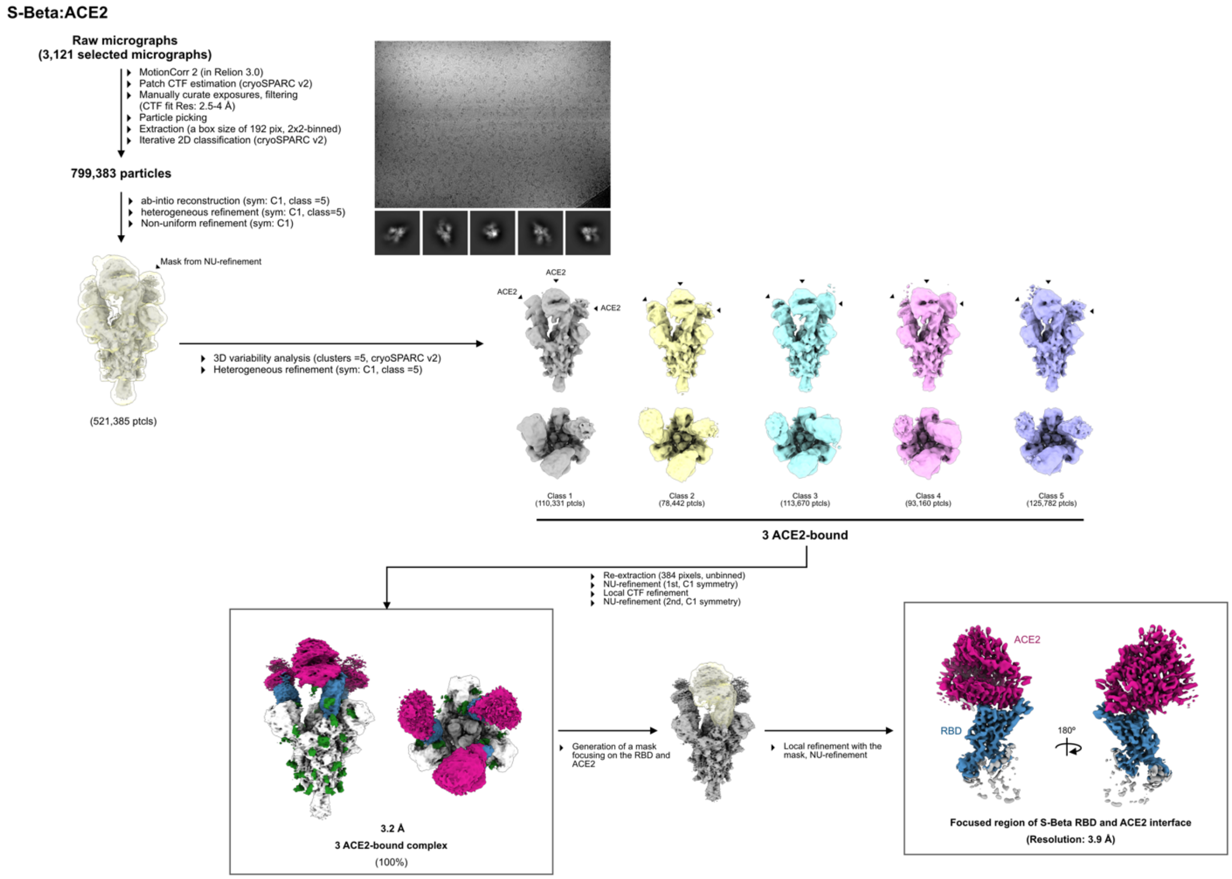

**Extended Data Fig. 8. Workflow for cryo-EM data processing of S-Beta in complex with ACE2.**

The same selection criteria of micrographs and processing procedure as described in *Extended Data Fig. 2* were used to yield five distinct structures after 3DVA and subsequent heterogeneous refinement. Particles corresponding to each 3D class were re-pooled and re-extracted after the structural inspection showing the similar conformation, yielding a final cryo-EM map of S-Beta:ACE2 with 3.2-Å resolution. To improve the resolution of ACE2-binding interface, a focused mask covering the RBD and ACE2 was used for the local refinement, generating a 3.9-Å cryo-EM map. ACE2 and RBD are colored in magenta and blue, respectively.

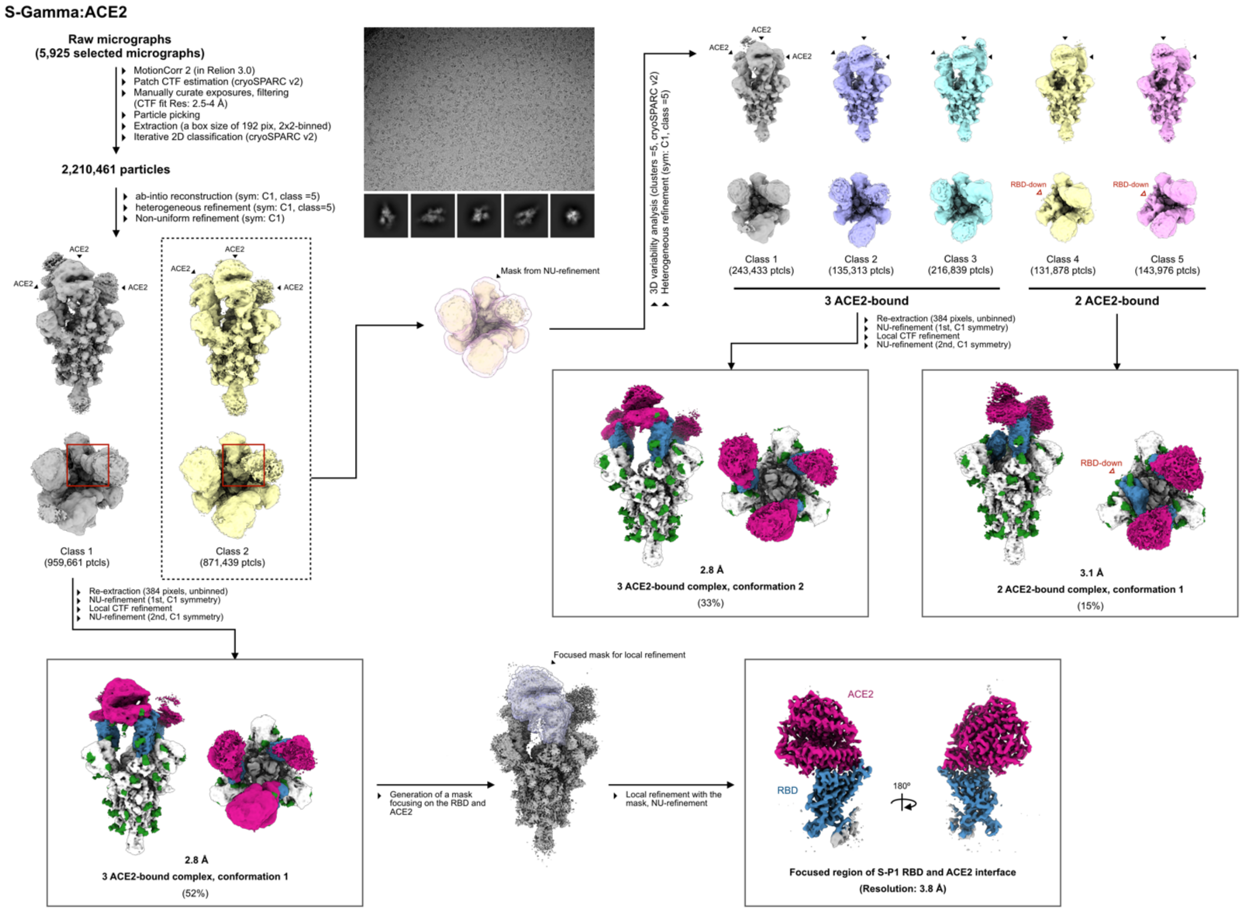

**Extended Data Fig. 9. Workflow for cryo-EM data processing of S-Gamma in complex with ACE2.**

Initial heterogeneous refinement followed by NU refinement yielded two 3 ACE-bound complexes, but one of two maps has a poorer resolution of the RBD region. Particles from these two 3D classes were therefore processing independently. Particles corresponding to the class with better resolution of RBD were directly refined by NU-refinement, yielding a 2.8-Å cryo-EM map with 3 ACE2-bound conformation. On the other hand, further 3DVA of the initial 3D class with poor RBD resolution demonstrated five classes, including three 3 ACE2-bound and two 2 ACE2-bound conformations. Particles from 3 ACE2-bound or 2 ACE2-bound conformations were subsequently re-pooled and re-extracted after the structural inspection showing similar conformation. A final round of NU refinement resulted in a 2.8-Å and a 3.1-Å cryo-EM maps in 3 ACE2-bound and ACE2-bound, respectively. The further local refinement focusing on the interface between RBD and

ACE was carried with the same strategy as described in *Extended Data Fig. 8* to generate a 3.8-Å cryo-EM map of this region.

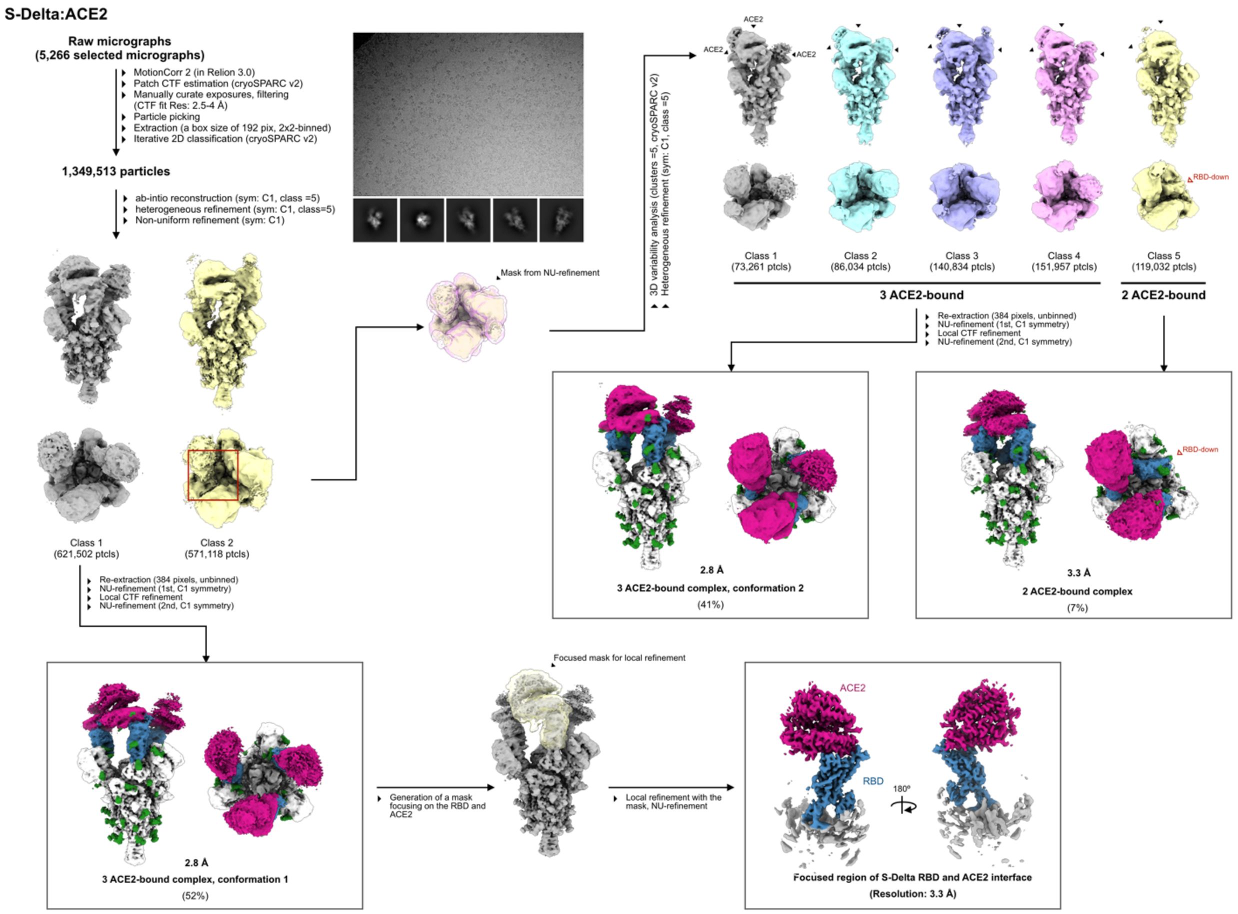

**Extended Data Fig. 10. Workflow for cryo-EM data processing of S-Delta in complex with ACE2.**

The same processing procedure as described in *Extended Data Fig. 9*. was used to yield three cryo-EM maps with 2.8 (3 ACE2-bound, conformation 1), 2.8 (3 ACE2-bound, conformation 2) and 3.3 Å (2 ACE2-bound), respectively. A further local refinement was implemented to improve the resolution of the ACE2-binding interface.

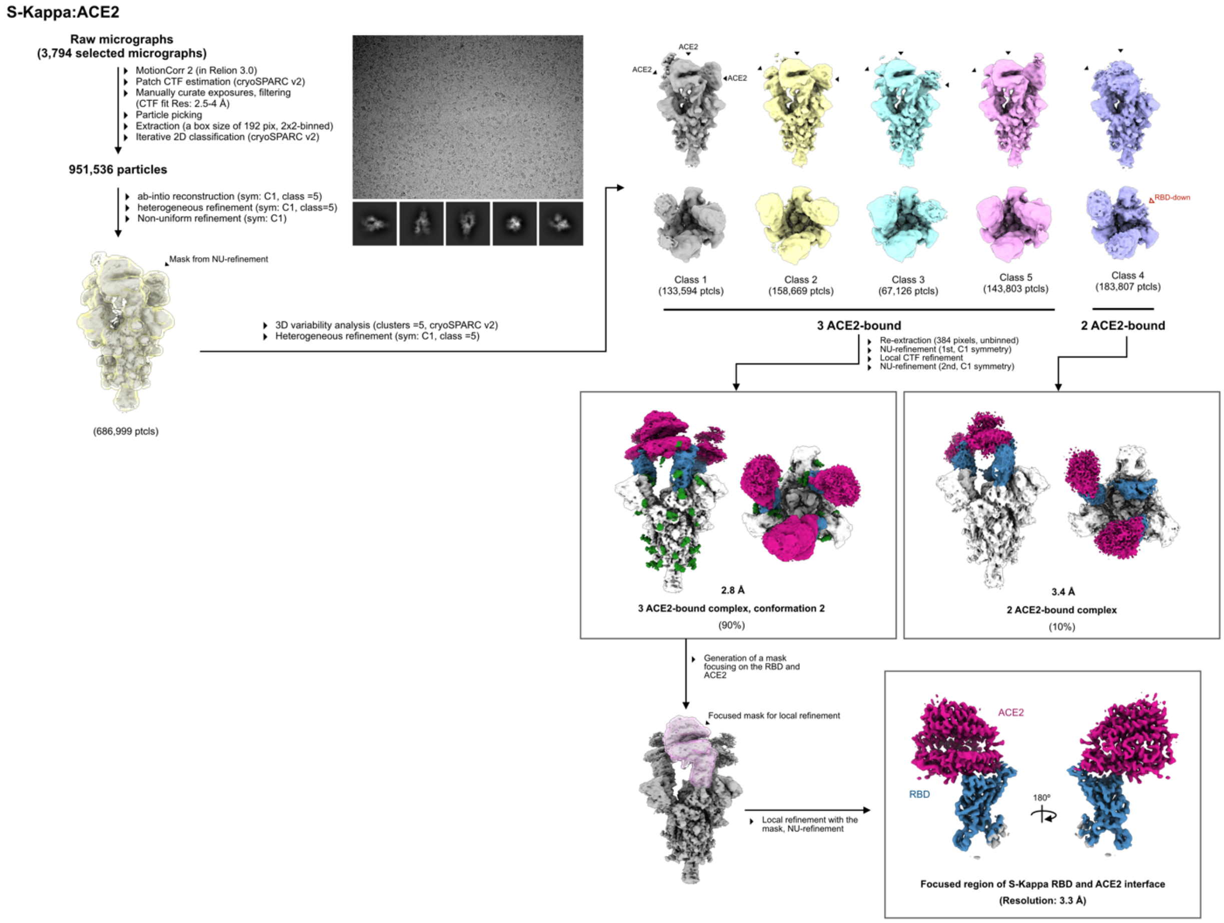

**Extended Data Fig. 11. Workflow for cryo-EM data processing of S-Kappa in complex with ACE2.**

The same processing procedure described in *Extended Data Fig. 9* yields four 3D classes based on the 3DVA. Particles from four 3 ACE2-bound classes were re-pooled and re-extracted due to the high structural similarity of these classes. Two independent NU-refinements were performed with two sets of particles, yielding one in 3 ACE2-bound conformation and one in 2 ACE2-bound conformation. The further local refinement was accomplished with a focused mask, generating a 3.3-Å cryo-EM to interpret the interface between RBD and ACE2 clearly.

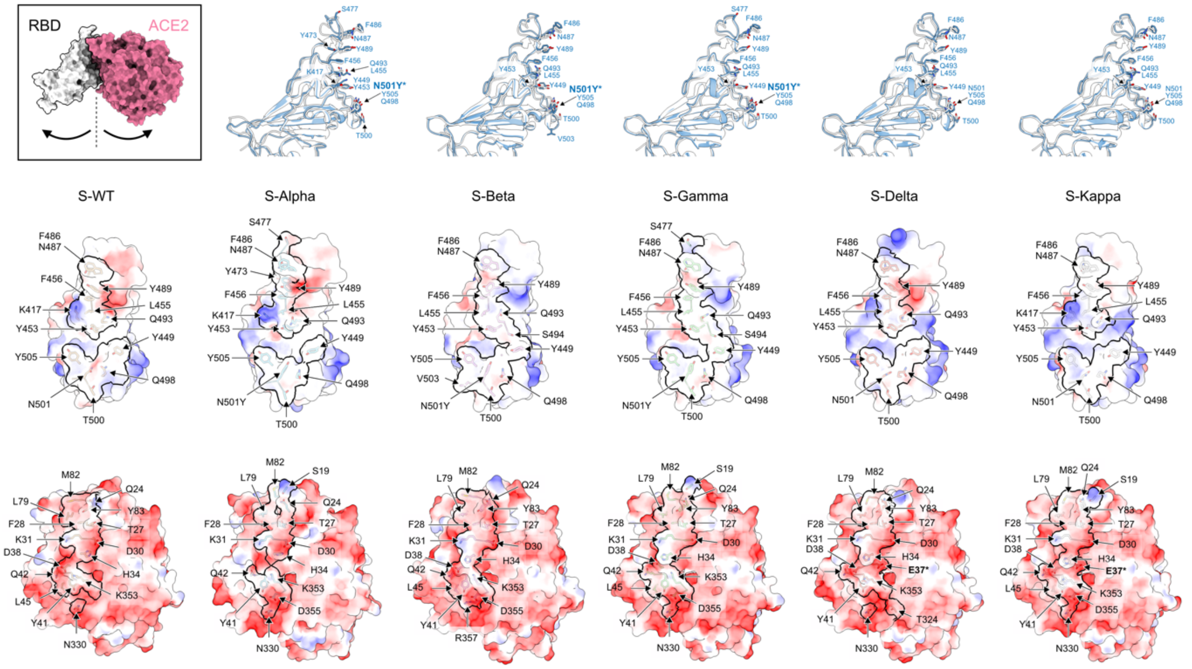

**Extended Data Fig. 12. Structural details of ACE2-binding interface of S variants.** (Top left inset) Surface representation of the pose of RBD-ACE2 interaction. (Top panel) Superpositions of the focus-refined RBD structures of individual S variants (blue) with that of S-WT (white, PDB ID: 6M0J). Residues that are involved in ACE2 binding are shown in cyan and white sticks for the variant and WT, respectively. Nitrogen and oxygen atoms are colored in blue and red, respectively. (Middle panel) Surface representations of the RBD of individual variants colored with the surface electrostatic potentials in blue and red for positive and negative charges, respectively. The ACE2 binding interface is outlined by black bold lines. Residues that are involved in ACE2 binding are shown in sticks with their identities indicated. (Bottom panel) Surface representations of ACE2 in complex with individual S variants following the same rendering scheme as used for RBD in the middle panel.

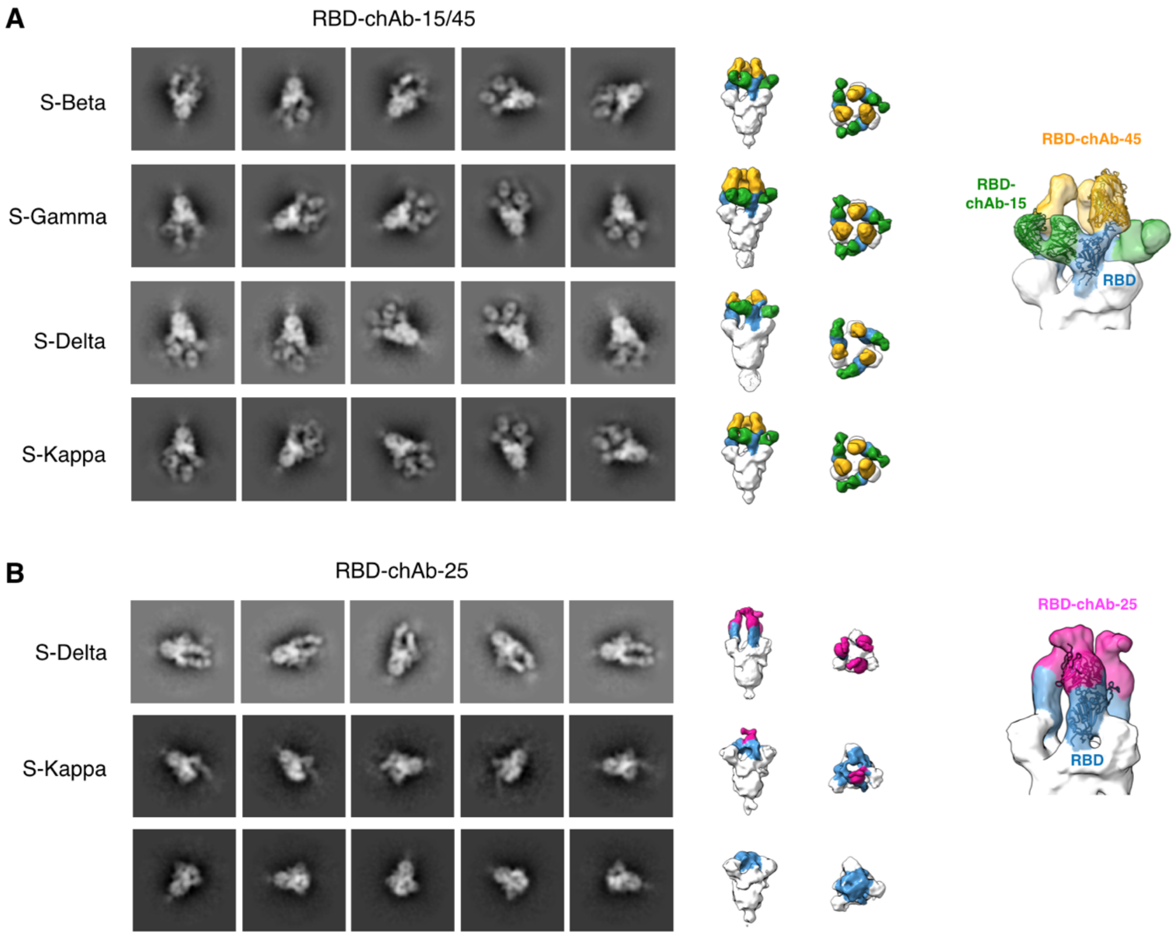

**Extended Data Fig. 13. 3D reconstruction of all S variants in complex with RBD-chAbs by negative staining electron microscopy (NSEM)**

A, Representative of 2D classification and 3D reconstruction of S-Beta, S-Gamma, S-Delta and S-Kappa in complex with RBD-chAb-15/45 cocktail. Side and top views of the final NSEM maps showing all S variants bind to RBD-chAb-15/45 in a 3:3:3 stoichiometry. RBD, RBD-chAb-15 and RBD-chAb-45 are colored in blue, green and yellow, respectively. (Top right) Superposition of the atomic models of S-RBD:RBD-chAb-15/45 complex (PDB ID: 7EH5) in the final NSEM map. B, Analysis of 2D classification and 3D reconstruction of S-Delta and S-Kappa in complex with RBD-chAb-25. The final NSEM map of S-Delta:RBD-chAb25 in the side and top view showing RBD-chAb-25 binds to S-Delta in a 3:3 stoichiometry. RBD-chAb-25 binds to S-Kappa primarily in an 1:3 stoichiometry. Additionally, a significant proportion of the particle images showed no nAb binding. The corresponding EM map indicated that the apo S-Kappa is likely in an all

RBD-down state. (Bottom right) Superposition of the atomic models of S-RBD:RBD-chAb-25 complex (PDB ID: 7EJ4) in the NSEM map.

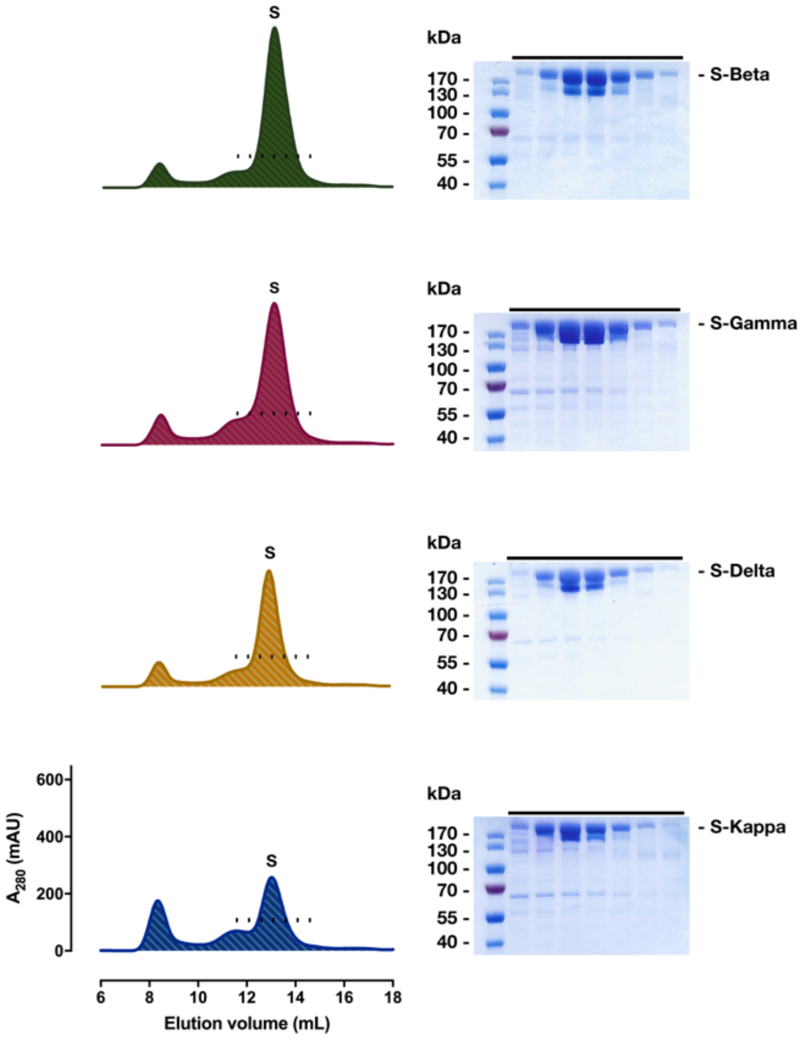

**Extended Data Fig. 14. Size exclusion chromatography and electrophoresis analysis of S variants.**

Size-exclusion chromatograms of S-Beta, S-Gamma, S-Delta and S-Kappa in descending order. The position of the main elution peak of each chromatogram that corresponds to the target S variant is indicated by "S" above. The positions of the individual fractions that were collected and analyzed by a 4-12% gradient SDS-PAGE (right panel) are indicated by dots. The SDS-PAGE gels were stained by Coomassie blue. The positions of the S variants are indicated individually on the right.

**Extended Data Table 1. Cryo-EM data collection, refinement and validation statistics for apo S-Beta (uncleavable form).**

| | 1 RBD-up conformation (EMD-31760) (PDB 7V76) | 2 RBD-up conformation (EMD-31761) (PDB 7V77) | 3 RBD-up conformation (EMD-31797) |
|---|---|---|---|
| **Data collection and processing** | | | |
| Magnification | 81000x | 81000x | 81000x |
| Voltage (kV) | 300 | 300 | 300 |
| Electron exposure ($e^{-}$/Å$^2$) | 50 | 50 | 50 |
| Defocus range (μm) | 0.8-2.6 | 0.8-2.6 | 0.8-2.6 |
| Pixel size (Å) | 1.1 | 1.1 | 1.1 |
| Symmetry imposed | C1 | C1 | C1 |
| Initial particle images (no.) | 2,667,093 | 2,667,093 | 2,667,093 |
| Final particle images (no.) | 306,270 | 289,915 | 288,721 |
| Map resolution (Å) | 3.2 | 3.3 | 3.5 |
| FSC threshold | 0.143 | 0.143 | 0.143 |
| **Refinement** | | | |
| Initial model used (PDB code) | 7EB3 | 7V76 (this study) | |
| Model resolution (Å) | 3.2 | 3.2 | |
| FSC threshold | 0.143 | 0.143 | |
| Map sharpening *B* factor (Å$^2$) | -104.2 | -100.7 | |
| Model composition | | | |
| Non-hydrogen atoms | 25,222 | 25,296 | |
| Protein residues | 3,088 | 3,102 | |
| Ligands | 77 | 77 | |
| *B* factors (Å$^2$) | | | |
| Protein | 32.11 | 42.83 | |
| Ligand | 72.84 | 81.12 | |
| R.m.s. deviations | | | |
| Bond lengths (Å) | 0.002 | 0.004 | |
| Bond angles (°) | 0.512 | 0.588 | |
| Validation | | | |
| MolProbity score | 1.65 | 1.87 | |
| Clashscore | 6.06 | 8.39 | |
| Poor rotamers (%) | 0.00 | 0.00 | |
| Ramachandran plot | | | |
| Favored (%) | 95.49 | 93.68 | |
| Allowed (%) | 4.47 | 6.22 | |
| Disallowed (%) | 0.03 | 0.10 | |

**Extended Data Table 2. Cryo-EM data collection, refinement and validation statistics for apo S-Beta (cleavable form).**

| | 1 RBD-up conformation (EMD-31798) (PDB 7V8C) | 2 RBD-up conformation (EMD-31799) | 3 RBD-up conformation (EMD-31800) |
|---|---|---|---|
| **Data collection and processing** | | | |
| Magnification | 81000x | 81000x | 81000x |
| Voltage (kV) | 300 | 300 | 300 |
| Electron exposure ($e^{-}$/Å$^2$) | 50 | 50 | 50 |
| Defocus range (μm) | 0.8-2.6 | 0.8-2.6 | 0.8-2.6 |
| Pixel size (Å) | 1.1 | 1.1 | 1.1 |
| Symmetry imposed | C1 | C1 | C3 |
| Initial particle images (no.) | 4,184,936 | 4,184,936 | 4,184,936 |
| Final particle images (no.) | 392,357 | 167,624 | 194,758 |
| Map resolution (Å) | 3.4 | 3.8 | 3.4 |
| FSC threshold | 0.143 | 0.143 | 0.143 |
| **Refinement** | | | |
| Initial model used (PDB code) | 7EB3 | | |
| Model resolution (Å) | 3.4 | | |
| FSC threshold | 0.143 | | |
| Map sharpening *B* factor (Å$^2$) | -104.1 | | |
| Model composition | | | |
| Non-hydrogen atoms | 25,200 | | |
| Protein residues | 3,088 | | |
| Ligands | 75 | | |
| *B* factors (Å$^2$) | | | |
| Protein | 64.07 | | |
| Ligand | 115.45 | | |
| R.m.s. deviations | | | |
| Bond lengths (Å) | 0.005 | | |
| Bond angles (°) | 0.614 | | |
| Validation | | | |
| MolProbity score | 1.99 | | |
| Clashscore | 10.69 | | |
| Poor rotamers (%) | 0.00 | | |
| Ramachandran plot | | | |
| Favored (%) | 93.03 | | |
| Allowed (%) | 6.81 | | |
| Disallowed (%) | 0.16 | | |

**Extended Data Table 3. Cryo-EM data collection, refinement and validation statistics for apo S-Gamma.**

| | 1 RBD-up conformation 1 (EMD-31762) (PDB 7V78) | 1 RBD-up conformation 2 (EMD-31763) (PDB 7V79) | 2 RBD-up conformation (EMD-31764) (PDB 7V7A) |
|---|---|---|---|
| **Data collection and processing** | | | |
| Magnification | 81000x | 81000x | 81000x |
| Voltage (kV) | 300 | 300 | 300 |
| Electron exposure ($e^{-}$/Å$^2$) | 50 | 50 | 50 |
| Defocus range (μm) | 0.8-2.6 | 0.8-2.6 | 0.8-2.6 |
| Pixel size (Å) | 1.1 | 1.1 | 1.1 |
| Symmetry imposed | C1 | C1 | C1 |
| Initial particle images (no.) | 3,129,900 | 3,129,900 | 3,129,900 |
| Final particle images (no.) | 327,212 | 535,097 | 346,258 |
| Map resolution (Å) | 3.4 | 3.3 | 3.4 |
| FSC threshold | 0.143 | 0.143 | 0.143 |
| **Refinement** | | | |
| Initial model used (PDB code) | 7EB3 | 7V78 (this study) | 7V78 (this study) |
| Model resolution (Å) | 3.3 | 3.2 | 3.4 |
| FSC threshold | 0.143 | 0.143 | 0.143 |
| Map sharpening *B* factor (Å$^2$) | -113.5 | -119.2 | -108.6 |
| Model composition | | | |
| Non-hydrogen atoms | 25,680 | 25,680 | 25,582 |
| Protein residues | 3,123 | 3,125 | 3,119 |
| Ligands | 90 | 89 | 86 |
| *B* factors (Å$^2$) | | | |
| Protein | 66.97 | 48.22 | 60.61 |
| Ligand | 105.26 | 80.64 | 100.45 |
| R.m.s. deviations | | | |
| Bond lengths (Å) | 0.004 | 0.005 | 0.002 |
| Bond angles (°) | 0.555 | 0.586 | 0.537 |
| Validation | | | |
| MolProbity score | 1.72 | 1.81 | 1.69 |
| Clashscore | 6.31 | 7.45 | 7.72 |
| Poor rotamers (%) | 0.00 | 0.00 | 0.04 |
| Ramachandran plot | | | |
| Favored (%) | 94.47 | 94.12 | 96.09 |
| Allowed (%) | 5.40 | 5.85 | 3.78 |
| Disallowed (%) | 0.13 | 0.03 | 0.13 |

**Extended Data Table 4. Cryo-EM data collection, refinement and validation statistics for apo S-Delta**

| | All RBD-down conformation (EMD-31775) (PDB 7V7N) | 1 RBD-up conformation 1 (EMD-31776) (PDB 7V7O) | 1 RBD-up conformation 2 (EMD-31777) (PDB 7V7P) |
|---|---|---|---|
| **Data collection and processing** | | | |
| Magnification | 81000x | 81000x | 81000x |
| Voltage (kV) | 300 | 300 | 300 |
| Electron exposure ($e^{-}$/Å$^2$) | 45.6 | 45.6 | 45.6 |
| Defocus range (μm) | 0.8-2.6 | 0.8-2.6 | 0.8-2.6 |
| Pixel size (Å) | 1.1 | 1.1 | 1.1 |
| Symmetry imposed | C1 | C1 | C1 |
| Initial particle images (no.) | 3,437,088 | 3,437,088 | 3,437,088 |
| Final particle images (no.) | 188,218 | 249,836 | 246,567 |
| Map resolution (Å) | 2.9 | 2.9 | 2.9 |
| FSC threshold | 0.143 | 0.143 | 0.143 |
| **Refinement** | | | |
| Initial model used (PDB code) | 7EB3 | 7V7N (this study) | 7V7O (this study) |
| Model resolution (Å) | 2.9 | 2.8 | 2.8 |
| FSC threshold | 0.143 | 0.143 | 0.143 |
| Map sharpening *B* factor (Å$^2$) | -79.7 | -83.3 | -84.7 |
| Model composition | | | |
| Non-hydrogen atoms | 25,433 | 25,328 | 25,374 |
| Protein residues | 3,101 | 3,092 | 3,096 |
| Ligands | 85 | 83 | 84 |
| *B* factors (Å$^2$) | | | |
| Protein | 48.88 | 40.54 | 43.44 |
| Ligand | 79.58 | 73.14 | 76.49 |
| R.m.s. deviations | | | |
| Bond lengths (Å) | 0.004 | 0.007 | 0.003 |
| Bond angles (°) | 0.547 | 0.630 | 0.545 |
| Validation | | | |
| MolProbity score | 1.74 | 1.76 | 1.60 |
| Clashscore | 6.60 | 6.95 | 5.52 |
| Poor rotamers (%) | 0.00 | 0.00 | 0.00 |
| Ramachandran plot | | | |
| Favored (%) | 94.43 | 94.45 | 95.64 |
| Allowed (%) | 5.44 | 5.49 | 4.20 |
| Disallowed (%) | 0.13 | 0.07 | 0.16 |

**Extended Data Table 4 (continue). Cryo-EM data collection, refinement and validation statistics for apo S-Delta.**

| | 1 RBD-up conformation 3 (EMD-31778) (PDB 7V7Q) | 1 RBD-up conformation 4 (EMD-31779) (PDB 7V7R) | 1 RBD-up conformation 5 (EMD-31780) (PDB 7V7S) |
|---|---|---|---|
| **Data collection and processing** | | | |
| Magnification | 81000x | 81000x | 81000x |
| Voltage (kV) | 300 | 300 | 300 |
| Electron exposure ($e^{-}/Å^2$) | 45.6 | 45.6 | 45.6 |
| Defocus range (μm) | 0.8-2.6 | 0.8-2.6 | 0.8-2.6 |
| Pixel size (Å) | 1.1 | 1.1 | 1.1 |
| Symmetry imposed | C1 | C1 | C1 |
| Initial particle images (no.) | 3,437,088 | 3,437,088 | 3,437,088 |
| Final particle images (no.) | 235,695 | 217,529 | 129,545 |
| Map resolution (Å) | 2.8 | 2.9 | 3.0 |
| FSC threshold | 0.143 | 0.143 | 0.143 |
| **Refinement** | | | |
| Initial model used (PDB code) | 7V7O (this study) | 7V7O (this study) | 7V7O (this study) |
| Model resolution (Å) | 2.8 | 2.8 | 3.0 |
| FSC threshold | 0.143 | 0.143 | 0.143 |
| Map sharpening *B* factor ($Å^2$) | -82.6 | -83.6 | -77.6 |
| Model composition | | | |
| Non-hydrogen atoms | 25,374 | 25,365 | 25,258 |
| Protein residues | 3,096 | 3,095 | 3,088 |
| Ligands | 84 | 84 | 81 |
| *B* factors ($Å^2$) | | | |
| Protein | 42.54 | 41.19 | 40.42 |
| Ligand | 72.95 | 74.81 | 71.25 |
| R.m.s. deviations | | | |
| Bond lengths (Å) | 0.003 | 0.003 | 0.003 |
| Bond angles (°) | 0.512 | 0.542 | 0.521 |
| Validation | | | |
| MolProbity score | 1.61 | 1.66 | 1.68 |
| Clashscore | 6.06 | 6.22 | 7.35 |
| Poor rotamers (%) | 0.00 | 0.00 | 0.00 |
| Ramachandran plot | | | |
| Favored (%) | 95.96 | 95.37 | 96.02 |
| Allowed (%) | 3.94 | 4.53 | 3.95 |
| Disallowed (%) | 0.10 | 0.10 | 0.03 |

**Extended Data Table 4 (continue). Cryo-EM data collection, refinement and validation statistics for apo S-Delta.**

| | 2 RBD-up conformation 1 (EMD-31781) (PDB 7V7T) | 2 RBD-up conformation 2 (EMD-31782) (PDB 7V7U) | 2 RBD-up conformation 3 (EMD-31783) (PDB 7V7V) |
|---|---|---|---|
| **Data collection and processing** | | | |
| Magnification | 81000x | 81000x | 81000x |
| Voltage (kV) | 300 | 300 | 300 |
| Electron exposure ($e^{-}/Å^2$) | 45.6 | 45.6 | 45.6 |
| Defocus range (μm) | 0.8-2.6 | 0.8-2.6 | 0.8-2.6 |
| Pixel size (Å) | 1.1 | 1.1 | 1.1 |
| Symmetry imposed | C1 | C1 | C1 |
| Initial particle images (no.) | 3,437,088 | 3,437,088 | 3,437,088 |
| Final particle images (no.) | 119,914 | 136,969 | 128,286 |
| Map resolution (Å) | 3.0 | 3.0 | 3.1 |
| FSC threshold | 0.143 | 0.143 | 0.143 |
| **Refinement** | | | |
| Initial model used (PDB code) | 7V7O (this study) | 7V7O (this study) | 7V7O (this study) |
| Model resolution (Å) | 3.0 | 3.0 | 3.0 |
| FSC threshold | 0.143 | 0.143 | 0.143 |
| Map sharpening *B* factor ($Å^2$) | -76.2 | -78.7 | -77.8 |
| Model composition | | | |
| Non-hydrogen atoms | 25,293 | 25,286 | 25,307 |
| Protein residues | 3,089 | 3,088 | 3,089 |
| Ligands | 83 | 83 | 84 |
| *B* factors ($Å^2$) | | | |
| Protein | 45.77 | 44.02 | 49.98 |
| Ligand | 78.16 | 78.23 | 82.61 |
| R.m.s. deviations | | | |
| Bond lengths (Å) | 0.003 | 0.005 | 0.002 |
| Bond angles (°) | 0.517 | 0.580 | 0.508 |
| Validation | | | |
| MolProbity score | 1.67 | 1.74 | 1.61 |
| Clashscore | 6.74 | 7.26 | 5.99 |
| Poor rotamers (%) | 0.00 | 0.00 | 0.00 |
| Ramachandran plot | | | |
| Favored (%) | 95.76 | 95.13 | 95.86 |
| Allowed (%) | 4.14 | 4.80 | 4.04 |
| Disallowed (%) | 0.10 | 0.07 | 0.10 |

**Extended Data Table 5. Cryo-EM data collection, refinement and validation statistics for apo S-Kappa.**

| | All RBD-down conformation (EMD-31767) (PDB 7V7D) | 1 RBD-up conformation 1 (EMD-31768) (PDB 7V7E) | 1 RBD-up conformation 2 (EMD-31769) (PDB 7V7F) | 2 RBD-up conformation (EMD-31770) (PDB 7V7G) |
|---|---|---|---|---|
| **Data collection and processing** | | | | |
| Magnification | 81000x | 81000x | 81000x | 81000x |
| Voltage (kV) | 300 | 300 | 300 | 300 |
| Electron exposure ($e^{-}$/Å$^2$) | 46 | 46 | 46 | 46 |
| Defocus range (μm) | 0.8-2.6 | 0.8-2.6 | 0.8-2.6 | 0.8-2.6 |
| Pixel size (Å) | 1.1 | 1.1 | 1.1 | 1.1 |
| Symmetry imposed | C1 | C1 | C1 | C1 |
| Initial particle images (no.) | 2,689,119 | 2,689,119 | 2,689,119 | 2,689,119 |
| Final particle images (no.) | 125,540 | 160,584 | 226,972 | 126,189 |
| Map resolution (Å) | 3.0 | 2.9 | 2.9 | 3.1 |
| FSC threshold | 0.143 | 0.143 | 0.143 | 0.143 |
| **Refinement** | | | | |
| Initial model used (PDB code) | 7EB3 | 7V7D (this study) | 7V7E (this study) | 7V7E (this study) |
| Model resolution (Å) | 3.0 | 2.9 | 2.9 | 3.0 |
| FSC threshold | 0.143 | 0.143 | 0.143 | 0.143 |
| Map sharpening *B* factor (Å$^2$) | -73.8 | -75.3 | -76.9 | -72.3 |
| Model composition | | | | |
| Non-hydrogen atoms | 25,356 | 25,343 | 25,506 | 25,464 |
| Protein residues | 3,095 | 3,093 | 3,114 | 3,114 |
| Ligands | 82 | 82 | 84 | 81 |
| *B* factors (Å$^2$) | | | | |
| Protein | 39.50 | 38.58 | 41.82 | 40.42 |
| Ligand | 76.11 | 71.49 | 78.70 | 78.66 |
| R.m.s. deviations | | | | |
| Bond lengths (Å) | 0.004 | 0.003 | 0.004 | 0.003 |
| Bond angles (°) | 0.564 | 0.553 | 0.561 | 0.554 |
| Validation | | | | |
| MolProbity score | 1.67 | 1.77 | 1.76 | 1.81 |
| Clashscore | 6.82 | 7.73 | 7.18 | 7.67 |
| Poor rotamers (%) | 0.00 | 0.00 | 0.00 | 0.04 |
| Ramachandran plot | | | | |
| Favored (%) | 95.70 | 94.98 | 94.75 | 94.32 |
| Allowed (%) | 4.27 | 4.96 | 5.12 | 5.61 |
| Disallowed (%) | 0.03 | 0.07 | 0.13 | 0.07 |

**Extended Data Table 6. Cryo-EM data collection, refinement and validation statistics for apo S-Kappa (dimer of S trimers).**

| | Conformation 1 (EMD-31771) (PDB 7V7H) | Conformation 2 (EMD-31772) (PDB 7V7I) | Conformation 3 (EMD-31773) (PDB 7V7J) |
|---|---|---|---|
| **Data collection and processing** | | | |
| Magnification | 81000x | 81000x | 81000x |
| Voltage (kV) | 300 | 300 | 300 |
| Electron exposure ($e^{-}/Å^2$) | 46 | 46 | 46 |
| Defocus range (μm) | 0.8-2.6 | 0.8-2.6 | 0.8-2.6 |
| Pixel size (Å) | 1.1 | 1.1 | 1.1 |
| Symmetry imposed | C1 | C1 | C1 |
| Initial particle images (no.) | 2,689,119 | 2,689,119 | 2,689,119 |
| Final particle images (no.) | 109,446 | 114,129 | 109,626 |
| Map resolution (Å) | 3.2 | 3.3 | 3.4 |
| FSC threshold | 0.143 | 0.143 | 0.143 |
| **Refinement** | | | |
| Initial model used (PDB code) | 7V7D (this study) | 7V7H (this study) | 7V7H (this study) |
| Model resolution (Å) | 3.2 | 3.3 | 3.4 |
| FSC threshold | 0.143 | 0.143 | 0.143 |
| Map sharpening *B* factor ($Å^2$) | -80.6 | -87.1 | -85.5 |
| Model composition | | | |
| Non-hydrogen atoms | 50,955 | 51,078 | 51,078 |
| Protein residues | 6,209 | 6,228 | 6,228 |
| Ligands | 174 | 174 | 174 |
| *B* factors ($Å^2$) | | | |
| Protein | 50.26 | 66.37 | 73.25 |
| Ligand | 133.48 | 142.65 | 142.33 |
| R.m.s. deviations | | | |
| Bond lengths (Å) | 0.002 | 0.005 | 0.004 |
| Bond angles (°) | 0.543 | 0.591 | 0.564 |
| Validation | | | |
| MolProbity score | 1.69 | 1.82 | 1.92 |
| Clashscore | 7.15 | 8.21 | 9.51 |
| Poor rotamers (%) | 0.00 | 0.00 | 0.00 |
| Ramachandran plot | | | |
| Favored (%) | 95.75 | 94.47 | 93.77 |
| Allowed (%) | 4.19 | 5.37 | 6.13 |
| Disallowed (%) | 0.07 | 0.16 | 0.10 |

**Extended Data Table 7. Cryo-EM and crystal sturctures used for NTD epitope counting**

| Antibody | PDB ID | EMDB | Reference |
|---|---|---|---|
| 4A8 | 7C2L | 30276 | [2] |
| FC05 | 7CWU | 30488 | [3] |
| 7M8J | CM25 | 23717 | [4] |
| 7NDD | COVOX-159 | 12284 | [5] |
| 7E8F | N9-Fab | 31017 | [6] |
| 7DZX | 8D2 | 30918 | [7] |
| 7DZY | 24900 | 30921 | [7] |
| 7NTC | P008-056 | 12587 | [8] |
| 7LXY | S2X333 | 23579 | [9] |
| 7LXZ | S2L28 | 23580 | [9] |
| 7LY2 | S2M28 | 23582 | [9] |
| 7L2E | 4-18 | 23126 | [10] |
| 7L2F | 5-24 | 23127 | [10] |
| 7L2D | 1-87 | 23125 | [10] |
| 7LAB | DH1052 | 23248 | [11] |
| 7LCN | D1050.1 | 23277 | [11] |
| 7N01 | 5-7 | 24097 | [12] |
| 7N62 | C12C9 | 24192 | [13] |
| 7N8I | S2L20 | 24237 | [14] |

**Extended Data Table 8. Kinetic parameters of ACE2 binding to spike variants derived from BLI.**

| | D614G[a] | Alpha[a] | Beta | Gamma | Delta | Kappa |
|---|---|---|---|---|---|---|
| $k_{on}$ ($10^4$ $s^{-1}$ $M^{-1}$) | 3.43 | 3.86 | 7.32 | 5.63 | 4.39 | 4.80 |
| $k_{off}$ ($10^{-5}$ $s^{-1}$) | 10.6 | 5.25 | 1.79 | 1.67 | 1.71 | 0.45 |
| $K_d$ ($10^{-9}$ M) | 3.10 | 1.36 | 0.25 | 0.30 | 0.39 | 0.09 |
| $K_d$ fold change[b] | 1 | 2.3 | 12.4 | 10.3 | 7.9 | 34.4 |

a. The values were taken from [15]

b. $K_d$ fold change is defined as $K_{d,D614G}/K_{d,variant}$

**Extended Data Table 9. Cryo-EM data collection, refinement and validation statistics for S-Beta in complex with ACE2.**

| | 3 ACE2-bound (EMD-31784) (PDB 7V7Z) | Local refinement of S-Beta-RBD:ACE2 (EMD-31785) (PDB 7V80) |
|---|---|---|
| **Data collection and processing** | | |
| Magnification | 81000x | 81000x |
| Voltage (kV) | 300 | 300 |
| Electron exposure ($e^{-}/Å^2$) | 50 | 50 |
| Defocus range (μm) | 0.8-2.6 | 0.8-2.6 |
| Pixel size (Å) | 1.1 | 1.1 |
| Symmetry imposed | C1 | C1 |
| Initial particle images (no.) | 2,400,152 | 2,400,152 |
| Final particle images (no.) | 518,601 | 518,601 |
| Map resolution (Å) | 3.1 | 3.9 |
| FSC threshold | 0.143 | 0.143 |
| **Refinement** | | |
| Initial model used (PDB code) | 7EDJ, 7V80 (this study) | 7EDJ |
| Model resolution (Å) | 3.1 | 3.8 |
| FSC threshold | 0.143 | 0.143 |
| Map sharpening *B* factor ($Å^2$) | -95.5 | -98.0 |
| Model composition | | |
| Non-hydrogen atoms | 39,857 | 6,610 |
| Protein residues | 4,884 | 797 |
| Ligands | 79 | 11 |
| *B* factors ($Å^2$) | | |
| Protein | 17.23 | 0.31 |
| Ligand | 55.35 | 4.92 |
| R.m.s. deviations | | |
| Bond lengths (Å) | 0.003 | 0.003 |
| Bond angles (°) | 0.573 | 0.614 |
| Validation | | |
| MolProbity score | 1.83 | 1.88 |
| Clashscore | 9.11 | 8.68 |
| Poor rotamers (%) | 0.00 | 0.00 |
| Ramachandran plot | | |
| Favored (%) | 95.07 | 93.82 |
| Allowed (%) | 4.93 | 6.18 |
| Disallowed (%) | 0.00 | 0.00 |

**Extended Data Table 10. Cryo-EM data collection, refinement and validation statistics for S-Gamma in complex with ACE2.**

| | 2 ACE2-bound (EMD-31786) (PDB 7V81) | 3 ACE2-bound Conformation 1 (EMD-31787) (PDB 7V82) | 3 ACE2-bound Conformation 2 (EMD-31788) (PDB 7V83) | Local refinement of S-Gamma-RBD:ACE2 (EMD-31789) (PDB 7V84) |
|---|---|---|---|---|
| **Data collection and processing** | | | | |
| Magnification | 81000x | 81000x | 81000x | 81000x |
| Voltage (kV) | 300 | 300 | 300 | 300 |
| Electron exposure ($e^{-}$/Å$^2$) | 50 | 50 | 50 | 50 |
| Defocus range (μm) | 0.8-2.6 | 0.8-2.6 | 0.8-2.6 | 0.8-2.6 |
| Pixel size (Å) | 1.1 | 1.1 | 1.1 | 1.1 |
| Symmetry imposed | C1 | C1 | C1 | C1 |
| Initial particle images (no.) | 4,710,579 | 4,710,579 | 4,710,579 | 4,710,579 |
| Final particle images (no.) | 274,310 | 591,860 | 953,630 | 953,630 |
| Map resolution (Å) | 3.2 | 2.8 | 2.8 | 3.0 |
| FSC threshold | 0.143 | 0.143 | 0.143 | 0.143 |
| **Refinement** | | | | |
| Initial model used (PDB code) | 7EDJ, 7V84 (this study) | 7V81 (this study) | 7V81 (this study) | 7EDJ |
| Model resolution (Å) | 3.1 | 2.8 | 2.8 | 2.9 |
| FSC threshold | 0.143 | 0.143 | 0.143 | 0.143 |
| Map sharpening *B* factor (Å$^2$) | -88.0 | -93.0 | -96.9 | -129.6 |
| Model composition | | | | |
| Non-hydrogen atoms | 35,252 | 40,098 | 40,168 | 6,637 |
| Protein residues | 4,311 | 4,902 | 4,902 | 797 |
| Ligands | 82 | 84 | 89 | 13 |
| *B* factors (Å$^2$) | | | | |
| Protein | 33.33 | 9.39 | 30.32 | 36.08 |
| Ligand | 66.95 | 56.12 | 61.95 | 65.05 |
| R.m.s. deviations | | | | |
| Bond lengths (Å) | 0.005 | 0.003 | 0.005 | 0.002 |
| Bond angles (°) | 0.620 | 0.558 | 0.582 | 0.493 |
| Validation | | | | |
| MolProbity score | 1.97 | 1.83 | 1.90 | 1.62 |
| Clashscore | 10.69 | 9.31 | 9.91 | 5.95 |
| Poor rotamers (%) | 0.03 | 0.00 | 0.00 | 0.00 |
| Ramachandran plot | | | | |
| Favored (%) | 93.47 | 95.17 | 94.31 | 95.71 |
| Allowed (%) | 6.50 | 4.83 | 5.69 | 4.29 |
| Disallowed (%) | 0.02 | 0.00 | 0.00 | 0.00 |

**Extended Data Table 11. Cryo-EM data collection, refinement and validation statistics for S-Delta in complex with ACE2.**

| | 2 ACE2-bound (EMD-31793) (PDB 7V88) | 3 ACE2-bound Conformation 1 (EMD-31794) (PDB 7V89) | 3 ACE2-bound Conformation 2 (EMD-31795) (PDB 7V8A) | Local refinement of S-Delta-RBD:ACE2 (EMD-31796) (PDB 7V8B) |
|---|---|---|---|---|
| **Data collection and processing** | | | | |
| Magnification | 81000x | 81000x | 81000x | 81000x |
| Voltage (kV) | 300 | 300 | 300 | 300 |
| Electron exposure ($e^{-}/Å^2$) | 46 | 46 | 46 | 46 |
| Defocus range (μm) | 0.8-2.6 | 0.8-2.6 | 0.8-2.6 | 0.8-2.6 |
| Pixel size (Å) | 1.1 | 1.1 | 1.1 | 1.1 |
| Symmetry imposed | C1 | C1 | C1 | C1 |
| Initial particle images (no.) | 3,456,891 | 3,456,891 | 3,456,891 | 3,456,891 |
| Final particle images (no.) | 85,521 | 481,781 | 617,671 | 617,671 |
| Map resolution (Å) | 3.3 | 2.8 | 2.7 | 3.2 |
| FSC threshold | 0.143 | 0.143 | 0.143 | 0.143 |
| **Refinement** | | | | |
| Initial model used (PDB code) | 7EDJ, 7V8B (this study) | 7V88 (this study) | 7V88 (this study) | 7EDJ |
| Model resolution (Å) | 3.2 | 2.8 | 2.7 | 3.2 |
| FSC threshold | 0.143 | 0.143 | 0.143 | 0.143 |
| Map sharpening *B* factor ($Å^2$) | -71.2 | -85.3 | -88.4 | -115.0 |
| Model composition | | | | |
| Non-hydrogen atoms | 35,103 | 40,040 | 39,928 | 6,651 |
| Protein residues | 4,281 | 4,869 | 4,869 | 797 |
| Ligands | 89 | 100 | 92 | 14 |
| *B* factors ($Å^2$) | | | | |
| Protein | 18.81 | 31.28 | 5.01 | 24.89 |
| Ligand | 65.53 | 55.15 | 52.54 | 55.53 |
| R.m.s. deviations | | | | |
| Bond lengths (Å) | 0.004 | 0.002 | 0.004 | 0.002 |
| Bond angles (°) | 0.581 | 0.524 | 0.528 | 0.529 |
| Validation | | | | |
| MolProbity score | 1.98 | 1.81 | 1.86 | 1.70 |
| Clashscore | 11.82 | 8.89 | 9.90 | 6.16 |
| Poor rotamers (%) | 0.00 | 0.00 | 0.00 | 0.00 |
| Ramachandran plot | | | | |
| Favored (%) | 94.25 | 95.16 | 95.06 | 94.83 |
| Allowed (%) | 5.68 | 4.80 | 4.92 | 5.04 |
| Disallowed (%) | 0.07 | 0.04 | 0.02 | 0.13 |

**Extended Data Table 12. Cryo-EM data collection, refinement and validation statistics for S-Kappa in complex with ACE2.**

| | 2 ACE2-bound (EMD-31790) (PDB 7V85) | 3 ACE2-bound (EMD-31791) (PDB 7V86) | Local refinement of S-Kappa-RBD:ACE2 (EMD-31792) (PDB 7V87) |
|---|---|---|---|
| **Data collection and processing** | | | |
| Magnification | 81000x | 81000x | 81000x |
| Voltage (kV) | 300 | 300 | 300 |
| Electron exposure ($e^{-}/Å^2$) | 45 | 45 | 45 |
| Defocus range (μm) | 0.8-2.6 | 0.8-2.6 | 0.8-2.6 |
| Pixel size (Å) | 1.1 | 1.1 | 1.1 |
| Symmetry imposed | C1 | C1 | C1 |
| Initial particle images (no.) | 2,786,914 | 2,786,914 | 2,786,914 |
| Final particle images (no.) | 66,721 | 616,121 | 616,121 |
| Map resolution (Å) | 3.3 | 2.8 | 3.3 |
| FSC threshold | 0.143 | 0.143 | 0.143 |
| **Refinement** | | | |
| Initial model used (PDB code) | 7EDJ, 7V87 (this study) | 7V85 (this study) | 7EDJ |
| Model resolution (Å) | 3.3 | 2.8 | 3.1 |
| FSC threshold | 0.143 | 0.143 | 0.143 |
| Map sharpening *B* factor ($Å^2$) | -60.4 | -86.8 | -108.3 |
| Model composition | | | |
| Non-hydrogen atoms | 35,106 | 40,002 | 6,618 |
| Protein residues | 4,306 | 4,881 | 797 |
| Ligands | 75 | 87 | 12 |
| *B* factors ($Å^2$) | | | |
| Protein | 37.09 | 15.06 | 11.33 |
| Ligand | 77.80 | 53.21 | 74.59 |
| R.m.s. deviations | | | |
| Bond lengths (Å) | 0.005 | 0.004 | 0.003 |
| Bond angles (°) | 0.624 | 0.555 | 0.588 |
| Validation | | | |
| MolProbity score | 2.06 | 1.86 | 1.72 |
| Clashscore | 12.69 | 9.24 | 6.35 |
| Poor rotamers (%) | 0.03 | 0.00 | 0.00 |
| Ramachandran plot | | | |
| Favored (%) | 93.16 | 94.66 | 94.58 |
| Allowed (%) | 6.75 | 5.30 | 5.42 |
| Disallowed (%) | 0.09 | 0.04 | 0.00 |

**Extended Data Table 13. NSEM data collection, refinement and validation statistics for VOC spikes in complex with nAbs**

| nAb | RBD-chAb-15/45 | | | | RBD-chAb-25 | |
|---|---|---|---|---|---|---|
| S protein | S-Beta (EMD-31817) | S-Gamma (EMD-31818) | S-Delta (EMD-31819) | S-Kappa (EMD-31820) | S-Delta (EMD-31821) | S-Kappa (EMD-31822) |
| **Data collection and processing** | | | | | | |
| Magnification | 50000x | 50000x | 50000x | 50000x | 50000x | 50000x |
| Voltage (kV) | 200 | 200 | 200 | 200 | 200 | 200 |
| Electron exposure (e–/Å$^2$) | 30 | 30 | 30 | 30 | 30 | 30 |
| Pixel size (Å) | 1.7 | 1.7 | 1.7 | 1.7 | 1.7 | 1.7 |
| Symmetry imposed | C3 | C3 | C3 | C3 | C3 | C1 |
| Initial particle images (no.) | 435,906 | 475,794 | 403,196 | 365,346 | 154,390 | 112,682 |
| Final particle images (no.) | 25,981 | 22,898 | 4,622 | 17,624 | 13,224 | 2,166 |
| Map resolution (Å) | 5.1 | 8.3 | 7.0 | 14.0 | 6.5 | 15.0 |
| FSC threshold | 0.143 | 0.143 | 0.143 | 0.143 | 0.143 | 0.143 |

**Extended Data Table 14. $IC_{50}$ values of nAbs against SARS-CoV-2 pseudovirus variants**

| nAb | $IC_{50}$ (ng/ml)[a] | | | | | |
|---|---|---|---|---|---|---|
| | D614G[b] | Alpha[b] | Beta | Gamma | Delta | Kappa |
| RBD-chAb-15 | 2.53 (1.64..3.89) | 36.4 (0.18..0.46) | 51.6 (25.6..109) | 6.03 (1.45..18.8) | 125 (94..173) | 123 (67..313) |
| RBD-chAb-25 | N.D. | N.D. | N.D. | N.D. | 47.7 (40.9..55.7) | 57.5 (33.6..108) |
| RBD-chAb-45 | 0.30 (0.18..0.46) | 3.67 (2.28..6.19)) | 3.46 (2.40..4.97) | 0.47 (0.29..0.72) | 20.1 (17.4..23.1) | 6.68 (5.35..8.31) |

a. The best-fit $IC_{50}$ values are shown above and the corresponding 95% confidence ranges are shown in parentheses below.
b. The values were taken from [15].

### References

1 Kuo, C.-W. *et al.* Distinct shifts in site-specific glycosylation pattern of SARS-CoV-2 spike proteins associated with arising mutations in the D614G and Alpha variants. *bioRxiv*, doi:10.1101/2021.07.21.453140 (2021).

2 Chi, X. *et al.* A neutralizing human antibody binds to the N-terminal domain of the Spike protein of SARS-CoV-2. *Science* **369**, 650-655, doi:10.1126/science.abc6952 (2020).

3 Wang, N. *et al.* Structure-based development of human antibody cocktails against SARS-CoV-2. *Cell Res* **31**, 101-103, doi:10.1038/s41422-020-00446-w (2021).

4 Voss, W. N. *et al.* Prevalent, protective, and convergent IgG recognition of SARS-CoV-2 non-RBD spike epitopes. *Science* **372**, 1108-1112, doi:10.1126/science.abg5268 (2021).

5 Dejnirattisai, W. *et al.* The antigenic anatomy of SARS-CoV-2 receptor binding domain. *Cell* **184**, 2183-2200 e2122, doi:10.1016/j.cell.2021.02.032 (2021).

6 Cao, Y. *et al.* Humoral immune response to circulating SARS-CoV-2 variants elicited by inactivated and RBD-subunit vaccines. *Cell Res* **31**, 732-741, doi:10.1038/s41422-021-00514-9 (2021).

7 Liu, Y. *et al.* An infectivity-enhancing site on the SARS-CoV-2 spike protein targeted by antibodies. *Cell* **184**, 3452-3466 e3418, doi:10.1016/j.cell.2021.05.032 (2021).

8 Rosa, A. *et al.* SARS-CoV-2 can recruit a heme metabolite to evade antibody immunity. *Sci Adv* **7**, doi:10.1126/sciadv.abg7607 (2021).

9 McCallum, M. *et al.* N-terminal domain antigenic mapping reveals a site of vulnerability for SARS-CoV-2. *Cell* **184**, 2332-2347 e2316, doi:10.1016/j.cell.2021.03.028 (2021).

10 Cerutti, G. *et al.* Potent SARS-CoV-2 neutralizing antibodies directed against spike N-terminal domain target a single supersite. *Cell Host Microbe* **29**, 819-833 e817, doi:10.1016/j.chom.2021.03.005 (2021).

11 Li, D. *et al.* In vitro and in vivo functions of SARS-CoV-2 infection-enhancing and neutralizing antibodies. *Cell* **184**, 4203-4219 e4232, doi:10.1016/j.cell.2021.06.021 (2021).

12 Cerutti, G. *et al.* Neutralizing antibody 5-7 defines a distinct site of vulnerability in SARS-CoV-2 spike N-terminal domain. *bioRxiv*, doi:10.1101/2021.06.29.450397 (2021).
13 Tong, P. *et al.* Memory B cell repertoire for recognition of evolving SARS-CoV-2 spike. *Cell*, doi:10.1016/j.cell.2021.07.025 (2021).
14 McCallum, M. *et al.* SARS-CoV-2 immune evasion by the B.1.427/B.1.429 variant of concern. *Science* **373**, 648-654, doi:10.1126/science.abi7994 (2021).
15 Yang, T. J. *et al.* Effect of SARS-CoV-2 B.1.1.7 mutations on spike protein structure and function. *Nat. Struct. Mol. Biol.*, doi:10.1038/s41594-021-00652-z (2021).

**Captions for Extended Movies**

**Extended Data Movie 1.**
Conformational landscape of SARS-CoV-2 S-Beta variant

**Extended Data Movies 2.**
Conformational landscape of SARS-CoV-2 S-Gamma variant

**Extended Data Movies 3**
Conformational landscape of SARS-CoV-2 S-Delta variant

**Extended Data Movies 4**
Conformational landscape of SARS-CoV-2 S-Kappa variant
